# The Cellular Expression and Genetics of Purple Body (*Pb*) in the Ocular Media of the Guppy *Poecilia reticulata*

**DOI:** 10.1101/121293

**Authors:** Alan S. Bias, Richard D. Squire

## Abstract

Our study revealed the presence of all major classes of chromatophores (melanophores, xanthophores, erythrophores, violet-blue iridophores, xantho-erythrophores) and crystalline platelets in various combinations in the iris and ocular media (cornea, aqueous humor, vitreous humor, outer lens membrane) of *Poecilia reticulata*. This novel ocular media study of *P. reticulata* takes into account the distinct interactions of Purple Body (*Pb*) based on results of previous Bias and Squire Purple Body (*Pb*) publications. Taken in conjunction with other researcher’s published results (regarding UV reflected color and pattern, vision, mate choice, individual preferences, and opsin capabilities) this indicates that these ocular chromatophore populations together create a complex ocular filter mechanism. This mechanism in turn provides spectral capabilities into the UV and Near-UV wavelengths in both Pb and non-Pb individuals. The chromatophores in the cornea, aqueous humor, covering membranes of the lens, and the vitreous humor comprise an ocular filter system that could reduce UV damage to the internal structures of the eye. The guppy’s ability to use UVA as a visual component provides a “private signally system” that cannot be detected by some predators. While non-Pb guppies should derive benefit in the near-UV from violet-blue iridophore units, greater benefit will be derived by Pb individuals with more violet iridophores functioning in the lower UV and near-UV wavelengths. To our knowledge little has been published for *P. reticulata* concerning pigmentation within the guppy eye. Macroscopic and microscopic imagery is presented.

---

**Fig 1.**
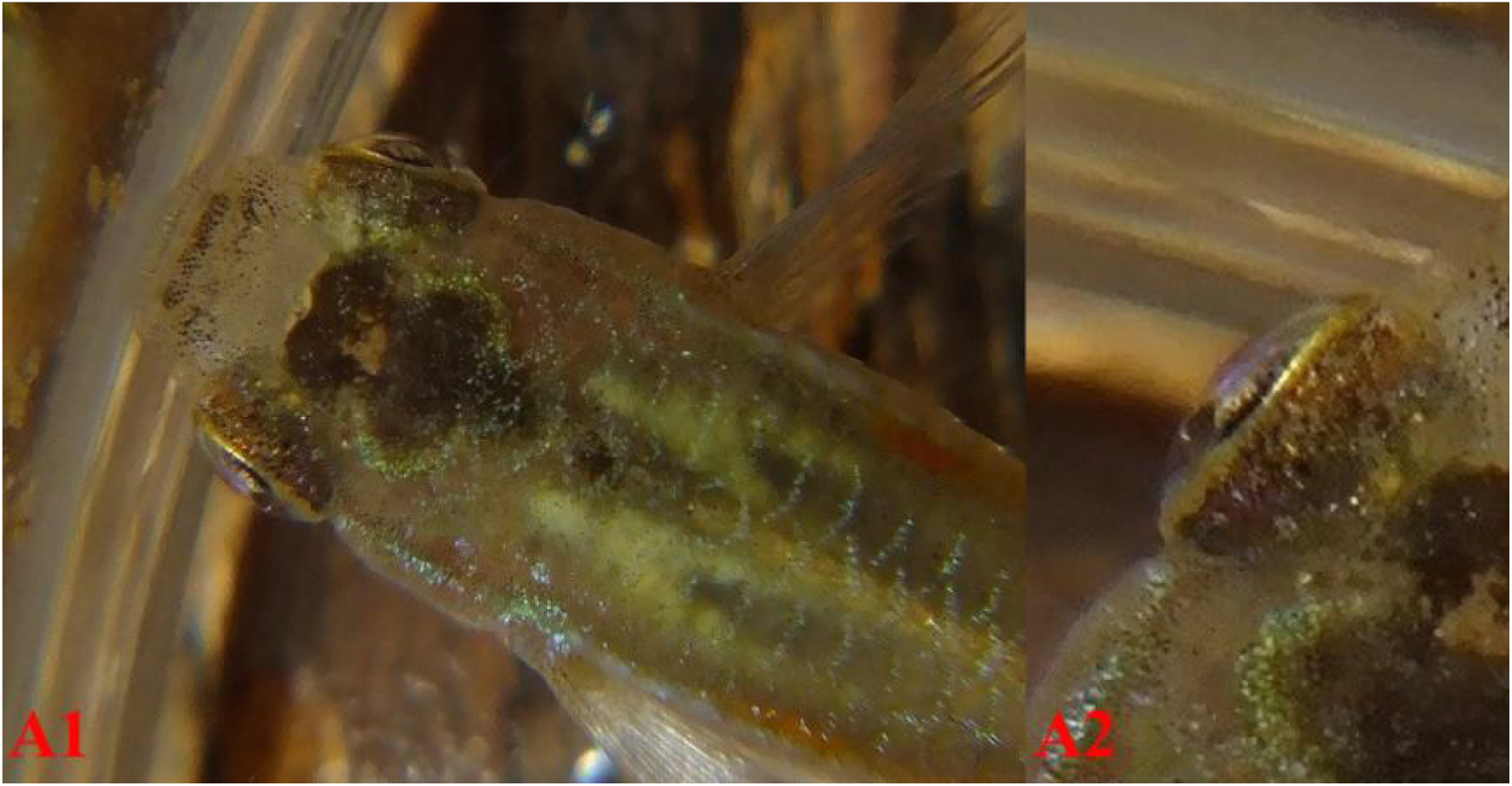
**(A1)** Jemez feral male Pb/-. **(A2)** Same field enlarged, high angle. Notice the protruding lens (PL) is reflecting ″violet″ from the iris in the central area of the pupil.

## Introduction

The intent of this paper is multifold: 1. To identify phenotypic and microscopic characteristics of the newly described Purple Body trait in ocular media. 2. To provide photographic and microscopic exhibits of Purple Body and non-Purple Body eyes for ease in identification of chromatophore types (Fig 2) and their interactions in the ocular media. 3. To encourage future study interest at a cellular level of populations in which Purple Body highlights UV (Ultra-Violet) and near-UV reflective qualities are found. 4. To stimulate molecular level studies of Purple Body and to identify the linkage group (LG) to which it belongs.

**Fig 2.**
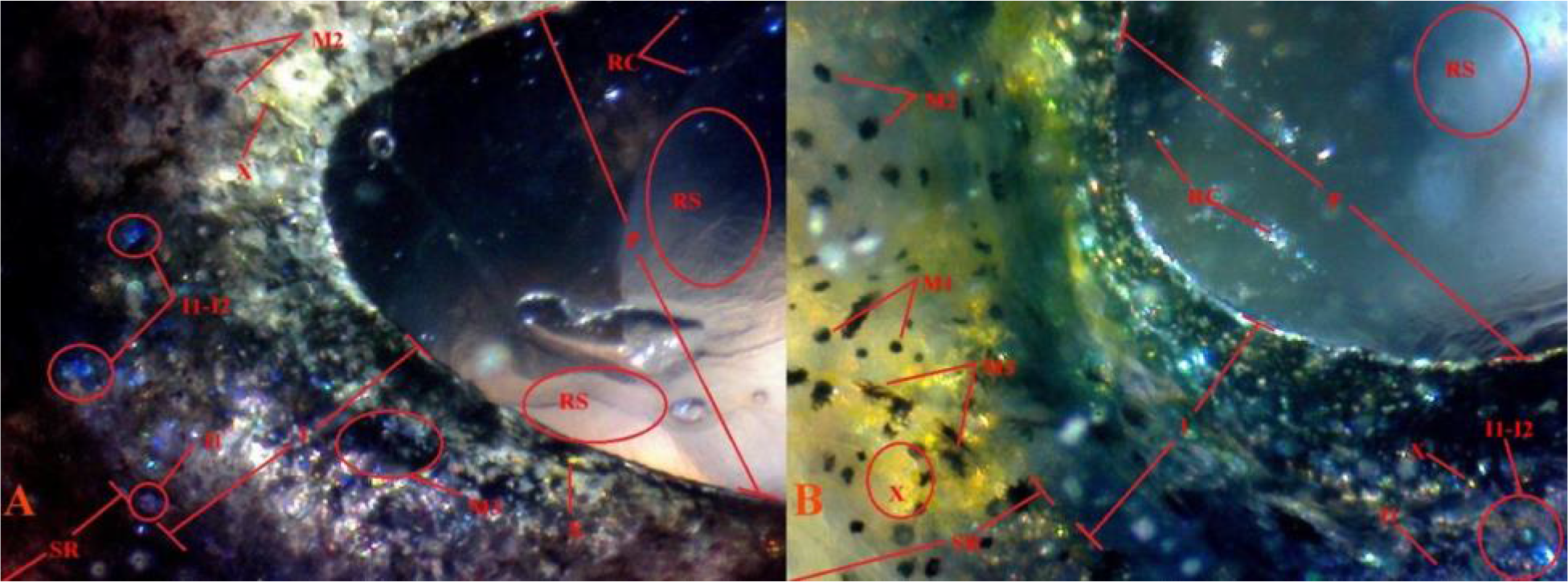
Pigment cell types and structures identified. **(A)** 17 Pb 40× 16 *Pb/pb (non-dissected pupil; iris and lens*) reflected light. **(B)** 22 non-Pb 40× 5 *pb/pb (dissected pupil; iris and lens*) reflected light. Melanophores punctate (M1), melanophores corolla (M2), melanophores dendritic (M3); to include visible dendritic melanophore strings and violet/blue iridophore chromatophore units. Violet iridophores (I1), blue iridophores (I2); to include violet-blue iridophore collections. Xanthophores (X); comprised of isolated single cells and small clustered groups (dendritic structures). Iris (I). Pupil (P). Scleral ring (SR). Reflected Chromatophore sheen from iris (RS). Reflected single cells from iris (RC).

Teleost species, including the Guppy, possess a complex eye with the ability to detect color and shape. Like many prey species, positioning of the eye is set for maximum field of view. Most species are considered to have fixed shape with adjustments made by changes in the amount of pupil protrusion; i.e. distance above the plane of the body. Variation in colors and color characteristics such as hue, depth, etc. cannot be important in female-based sexual selection unless the female, and male, can detect these color characteristics. Therefore, the evolution of color characteristics must be accompanied by the evolution of the ability to detect these colors. Endler showed that selection for spectral sensitivity variation in both short-wavelength sensitivity (SWS) and long wave sensitivity (*LWS*) is due to a hereditable factor in guppies (Endler 2001).

To our knowledge little has been published for *P. reticulata* concerning pigmentation within the guppy eye (Kunz 1977). Recent study indicates color vision varies across populations, and that populations with stronger preferences for orange had higher LWS opsin levels (Sandkam 2015a **and** 2015b). It has been shown that Guppies are able to perceive UV wavelengths, and that males reflect UV from both structural color and color pigment with variability between individuals. It was further shown that female association preference with males occurs under long wavelength (*UV-A*) conditions in which orange is visible (White 2003). While this study suggested that females have little or no sexual selective preference for either low UV or high UV males, it did not specifically focus on any benefit derived from reflective qualities of Pb under reduced ambient lighting conditions.

Early studies re-affirmed that courtship activity was at its highest during dawn and dusk. These are periods during which SWS and LWS are visible from low angle ambient sunlight (Endler 1987, 1991, 1992; Loew 1990). Others confirm the presence of UV-sensitive retinal cones, UV-transmittable ocular media, and SWS opsin genes in guppies (Douglas 1989, 1990; Archer 1987, 1990; Weadick 2007; Ward 2008; Watson 2010; Smith 2002). Individuals expressing Pb exhibit higher violet to blue iridophore density. Whether this results from an increased number of cells or simply the result of their increased visibility from the reduction of yellow xanthophores has not been determined. Existing red erythrophore populations appear unaltered (Bias and Squire, 2017a).

While Archer (1987) was unable to prove the existence of visual pigments extending into the accepted starting range for peak sensitivity (*maximum absorbance* - λ_max_) in UV spectrum (UVA *380-400nm*), he showed well marked clusters at λ_max_ 410nm, 465nm and 573nm. He concurred with earlier studies asserting that Guppies are polymorphic for color vision in LWS, with most rhodopsin-porphyropsin polymorphism in cones absorbing yellow, orange and red. Kemp in turn reported UV reflectance of violet-blue iridophores and orange spots ranging from 350-400nm (Kemp 2008). A molecular level study (Ward 2008) indicates a higher than normal duplication and divergence of 4 distinct LWS in *Poecilia*, as compared to other species. Laver (2011) conducted PCR studies that showed the presence of 11 different opsin genes in guppies originating from Cumaná, Venezuela. They found that 10 different opsins are found in juveniles, and both male and female adults.

With the discovery of variation in opsin expression between individuals of the Guppy’s eleven opsin cones, it has been suggested that new designs in behavioral study are warranted in regard to mate choice (Rennison 2011). Modification of scleral and iris pigment is noted in Pb, resulting in greatly increased levels of violet iridophores as compared to non-Pb. A similar situation is also found with modification by other traits, such as Metal Gold (Mg) (Bias 2015, and *unpublished breeding notes*), that produces not only proliferation of reflective yellow color pigments in the body, but also in ocular media (cornea, aqueous humor, vitreous humor, outer lens membrane).

## Materials

### ID Number, Pb or non-Pb, Color / Strain, Genotype

**(*See:*** Supplemental S1 for Strain Genotypes and Slide Specimen Photos).

13 Pb male (grey E) *Pb/Pb*.
17 Pb (grey E, litter mate – not actual male) *Pb/pb*.
18 Pb (grey E, related male – not actual male) *Pb/Pb*.
19 non-Pb (grey E, litter mate – not actual male) *pb/pb*.
22 non-Pb (grey) *pb/pb*.
23 Pb (grey) *Pb/pb*.
28 non-Pb (blond Ginga) *pb/pb*.
30 Pb [Jemez Female] (grey) *Pb/-*.
31 Pb [Jemez Female] (grey) *Pb/-*.
32 Pb [Jemez Female] (grey) *Pb/-*.
33 Pb [Jemez Female] (grey) *Pb/-*.
34 Pb [Jemez Female] (grey) *Pb/-*.

## Methods

All study fish were raised in 5.75, 8.75 and 10-gallon all-glass aquaria dependent upon age. They received 16 hours of light and 8 hours of darkness per day. Temperatures ranged from 78°F to 82°F. Fish were fed a blend of commercially available vegetable and algae based flake foods and Ziegler Finfish Starter (50/50 mix ratio) twice daily, and newly hatched live Artemia nauplii twice daily. A high volume feeding schedule was maintained in an attempt to produce two positive results: 1. Reduce the time to onset of initial sexual maturity and coloration, thus reduce time between breedings. 2. Increase mature size for ease of phenotypic evaluation and related microscopic study.

All euthanized specimens were photographed immediately, or as soon as possible, after temperature reduction (rapid chilling) in water (H_2_0) at temperatures just above freezing (0°C) to avoid potential damage to tissue and chromatophores, while preserving maximum expression of motile xantho-erythrophores in Pb and non-Pb specimens. All anesthetized specimens were photographed immediately after short-term immersion in a mixture of 50% aged tank water (H20) and 50% carbonated water (H_2_CO^3^).

All dried specimens were photographed immediately after rehydration in cold water (H_2_0). Prior euthanasia was by cold water (H_2_0) immersion at temperatures just above freezing (0 °C). MS-222 (Tricaine methanesulfonate) was not used to avoid the potential for reported damage and/or alterations to chromatophores, in particular melanophores, prior to slide preparation.

## Results

### I. Description and Characteristics: Pb (*Pb/Pb*) vs. non-Pb (*pb/pb*)

Our results show chromatophore populations residing in all areas of ocular media with the possible exception of the lens itself. We postulate that dense layers of violet-blue iridophores in conjunction with melanophores and xanthophores residing within the cornea-aqueous humor-iris-vitreous humor and the surrounding capsule at the anterior pole of the crystalline lens act as “ocular media filters”, with individuals deriving benefit in the UV and/or near-UV spectrum. The existence of similar filters has been described and summarized in other teleost fish species and mammals (Douglas 1989, 1999, 2014; Siebeck 2001). Pb will provide benefit at lower wavelengths with increased levels of violet iridophores, and non-Pb will have reduced benefit at slightly higher average wavelengths with balanced violet-blue iridophores. Xanthophores in turn, counter balance and provide benefit in the higher wavelengths.

Douglas states, “The range of wavelengths to which an animal is sensitive depends both on the spectral location of its visual pigments and on the wavelengths that impinge upon them. The latter is governed not only by the environment in which the animal lives but also by the absorption and reflection characteristics of structures within the eye. Thus, any consideration of fish colour vision must take into account the transmission of their lens and cornea” (Douglas 1989). Prior to this, ocular media was commonly considered to be “clear” for the most part in both freshwater and marine species. The exceptions were species noted with yellow corneas (see Douglas 1989 for review). Advances in conventional microscopy through the use of digital cameras and software allow us to clearly show the presence of structural and pigmented color residing in locations that earlier appeared to be clear under transmitted light.

Spectral color is produced by single wavelengths of ambient sunlight. The human Visible Wave Length Band (Visual Color) includes: red (*620-670 nm* Bright Red / *670-750 nm* Dark Red), orange and yellow (*570-620 nm*), green (*500-570 nm*), blue (*430-500 nm*), and violet (*400-430 nm*). Red light, with the longest wavelength and the least amount of energy, allows natural light penetration at less depth. Blue / violet light (*near-UV*), has the shortest wavelength and the most amount of energy, and allows natural light penetration to greater depth. Violet is a true wavelength color, while Purple is a composite effect produced by combining blue and red wavelength colors. Since we as humans automatically think of the visual spectrum in terms of what we ourselves can see, it is all too easy to forget that for guppies, the visual spectrum extends down into the UVA range of at least 250-400nm (see the Discussion for more details and references).

In general, while there are microscopic differences, our findings of visual distinctions between Pb and non-Pb are often more consistent, as opposed to microscopic distinctions. Much of this is likely the result of variability in both zygosity and ornament composition between individuals, within and between both populations and strains. Microscopically, structural differentiation between xantho-erythrophores appears minimal, with differences in the population levels and collection or clustering of xanthophores. Heterozygous Pb exhibits partial reduction in collected xanthophores, and homozygous Pb exhibits the near complete removal of collected and clustered xanthophores. Though, it is noted that yellow color cell populations consisting of isolated “wild-type” single cell xanthophores remain intact.

### II. Macroscopic Observations: The eye of *Poecilia reticulata* in homozygous Pb *Pb/Pb*, heterozygous Pb *Pb/Pb* and non-Pb *Pb/Pb*

Our results are based on several phenotypes and multiple specimens. The cornea and pupil are circular shaped allowing for a near 360° wide angle view of their environment, with little bending of light wavelengths during transmission. Light adjustment is accomplished primarily by dorsoventral and anteroposterior adjustment of the iris. In the iris Pb exhibited an “overall” higher incidence of violet iridophores and “purple” appearance, as compared to non-Pb that express a “blue appearance with either a predominance of blue iridophores or equal ratio of violet-blue iridophores. ***[Note:*** *hereafter referenced as balanced to reflect a predominance of blue or equal violet-blue iridophores for ease of discussion*].

*P. reticulata* cranial structure is bilaterally symmetric when viewed from a high angle. The Left-right axis gently slopes from the dorsal base in even taper to the supraocciptal surface (see S1 for naming and locations of axial planes). Then a slight increase in taper begins and continues to the mouth (Fig 3A-C). Differential between males and females is minimal, though greater between individuals.

**Fig 3.**
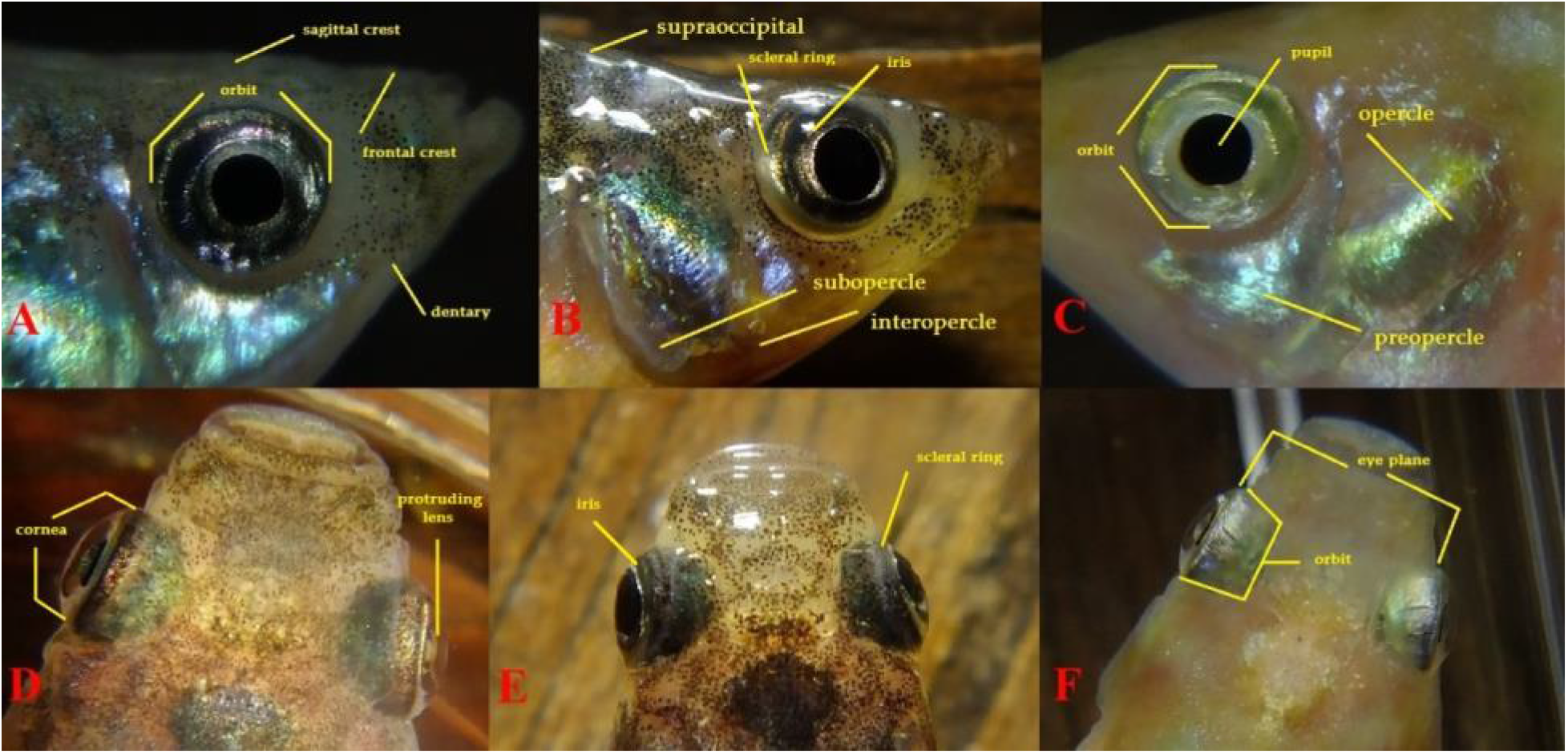
**(A)** Grey dark-eye dominant **Pb/Pb** female. **(B)** Grey Light Eye Pb/- female. **(C)** Grey Non-Pb **Pb/Pb** female. Chromatophores (violet-blue iridophores, melanophores and xanthophores) are visible in both the scleral ring and iris of all specimens. The cornea and underlying aqueous humor fluid appear clear to the naked eye above the proximal pupil region.

The dorsoventral axis is also generally bilaterally symmetric. Operculum (gill plate) is observed to consist of fused bony opercle, preopercle, interopercle and flexible subopercle. Dorsal side slopes downward starting at the dorsal base, increasingly past the supraocciptal to the upper jaw. The ventral side axis maintains a more general upward slope to the subopercle, with increasing upward angle past the interopercle through the dentary to the lower jaw (Fig 3 **and** 4, D-F). Differential between males and females is minimal, though greater between individuals and often appears more consistent among males

**Fig 4.**
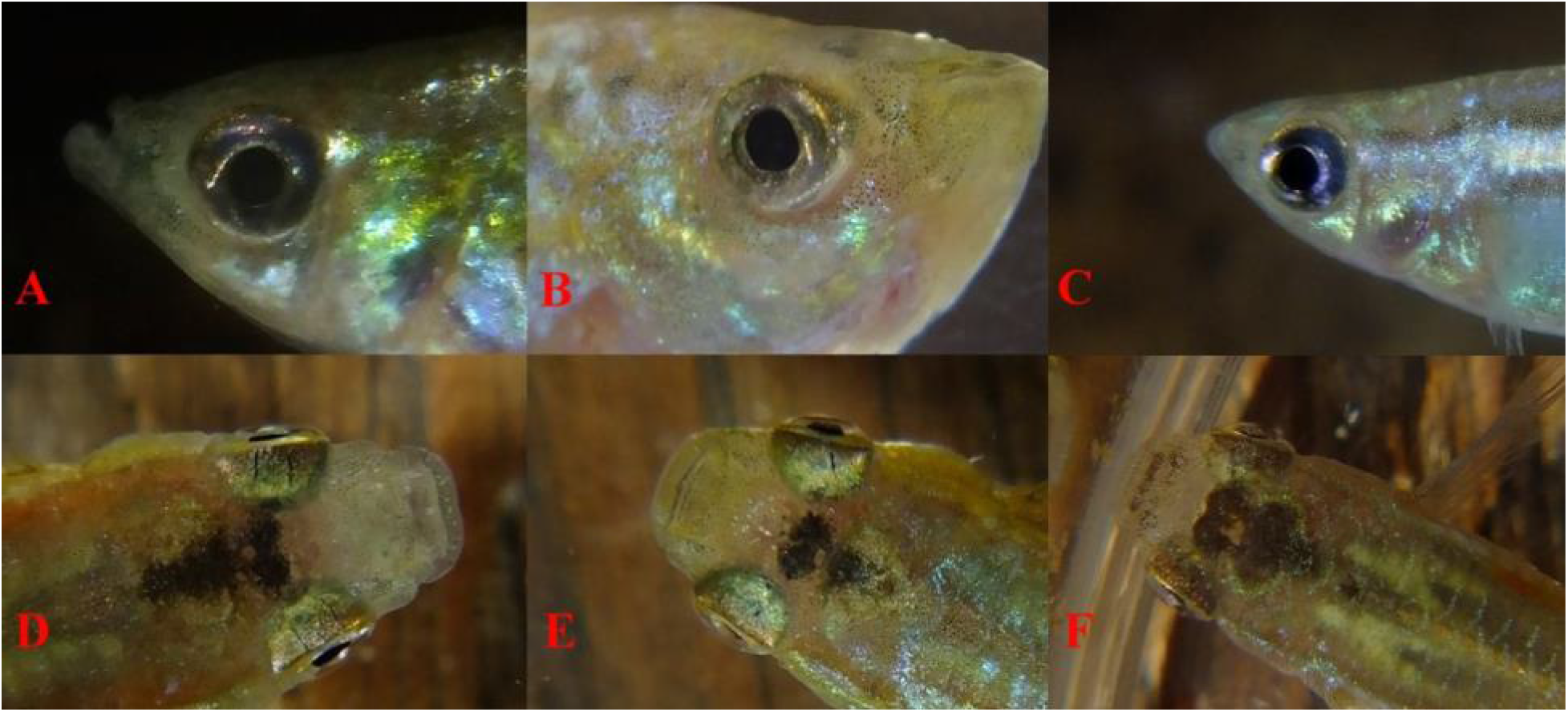
**(A & D)** Grey **Pb/Pb** male. **(B & E)** Grey Pb/- male. **(C & F)** Grey Non-Pb **Pb/Pb** male. Chromatophores (violet-blue iridophores, melanophores and xanthophores) are visible in both the scleral ring and iris of all specimens. Cornea and underlying aqueous humor fluid appear clear to the naked eye above the proximal pupil region.

High-angle macroscopic images reveal lens protrusion well past the plane of the iris to produce a wide field of view (Fig 3 **and** 4, D-F), with a corneal size near equal to the circumference of the eye extending to the scleral ring. Eyes are deep set with high chromatophore content in bony orbits between the sagittal crest and the preopercle. The overall forward pointing of the eye-set generally follows the tapering of the left-right axis. Variability between males and females is minimal, though it often appeared greater between females and more consistent among males. Pupils express no visible aphakic gap (the “lensless” part of the pupil that does not cover the lens, Schmitz 2011) between the protruding lens and iris in perpendicular macroscopic images (Fig 3 **and** 4, A-C). This would be expected in the case of freshwater herbivore/detritivore prey species (Gagnon 2016).

The pupillary response to ambient light changes in bony teleosts is generally considered static; i.e. non-responsive. Limited research has shown several species are capable of pupil dilation (Douglas 1998; Schmitz 2011). *Poecilia reticulata* populations and Domestic strains commonly possess two distinct eye types: “black or dark-eye” (Fig 3A-B, 4A, 4C) and “silver or light eye” (Fig 3C, 4B), based on coloration of the sclera and iris. In *P. reticulata* dark eye is often associated with dominance or aggression and the light eye are more prevalent. This is in contrast to some species of cichlidae (Miyai 2011). In some *P. reticulata* populations and strains a portion of individuals may remain consistently dark-eyed, while in others only dominant male(s) and female(s) express dark-eyes (Gorlick 1976; Martin 1981; Magurran 1991).

This demonstrates an ability for pupillary response in the form of dilation in *P. reticulata*, though not necessarily to changes in ambient lighting. Regardless of eye type, dense populations of epithelial violet-blue iridophores and melanophores were observed macroscopically in the iris in Pb phenotypes producing a more “purple” appearance. In contrast, non-Pb irises tended to express balanced violet-blue iridophore and equal melanophore populations in both eye types with “blue” appearance. Again, there was much variability in observations and the complete genotype and angle of observation for each individual specimen had to be considered in interpretation and understanding of the results.

Observations of the visual axis in the Guppy reveal the eye tilts at a downward angle (Fig 5A) and slightly forward (Fig 5B) from the tapering body structure. Other than occasional “reflex blinking” to adjust the iris and / or lens, movements that are common in teleost species without the benefit of an eyelid, the eyes are static. This reflex movement is assumed to be a mechanism for muscle relaxation and/or refocusing of the eye. Convex shape and curvature in the plane of the iris was detected in high-angle macroscopic images.

**Fig 5.**
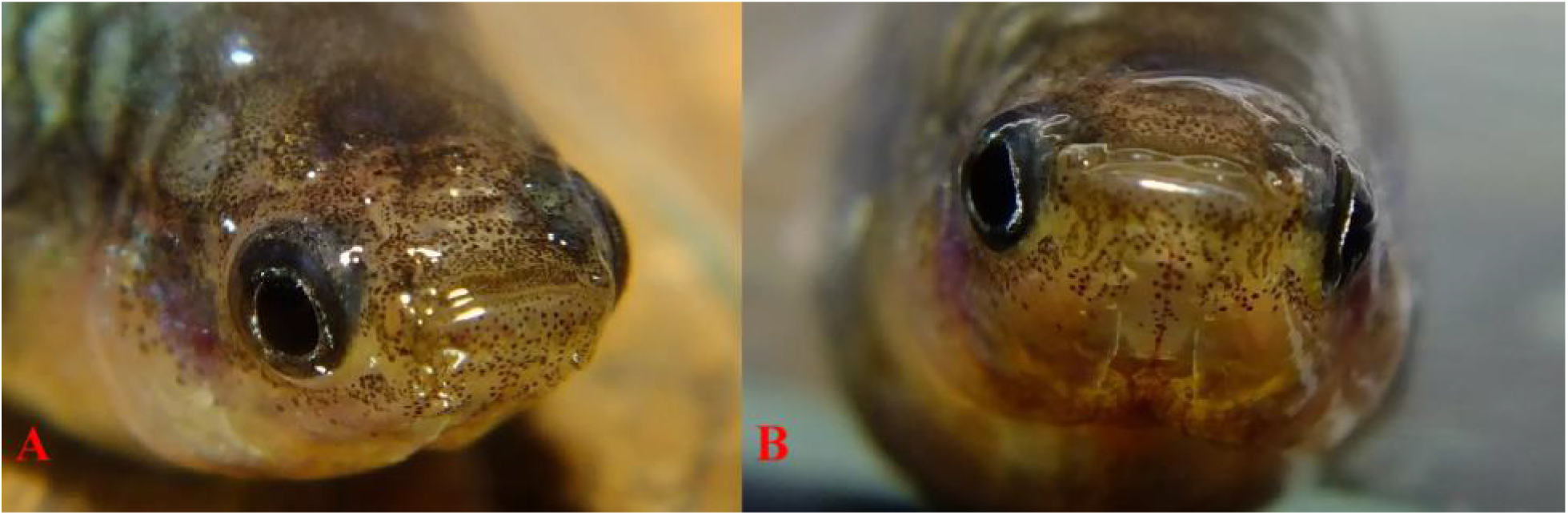
**(A-B)** Jemez feral female (Pb/-). Live, anesthetized. This female was removed from a breeding group and is expressing dominant ″dark-eye″, though this feature is often expressed in multiple females in this population when housed together. Both the iris and scleral ring contain melanophores, violet-blue iridophores and xanthophores.

Visual observations, in two forms, of live specimens and photographic images indicate that lack of eye movement is somewhat compensated for by control of lens movement in angle and direction (Fernald 1988; Gagnon 2016). First, observations at a perpendicular angle show the complete circular nature of the pupillary shape with visible overlap from the iris and lack of aphakic gap correlated with high angle observations showing tilting of the lens. Second, differences in width between the anterior and posterior iris was often observed, indicating directional control of the protruding lens. Further research is needed to determine capabilities and limitations of multifocal lens in the Guppy.

### III. Microscopic Observations: The eye of *Poecilia reticulata* in homozygous Pb *Pb/Pb*, heterozygous Pb *Pb/Pb* and non-Pb *Pb/Pb*

The ocular media of *P. reticulata* is comprised of a cornea, aqueous humor fluid, iris, lens, and surrounding vitreous humor fluid through which light passes to the retina. The lens of the Guppy is spherical in shape, allowing for a high degree of light refraction. Vision is clearest in the central portion of the eye and weakest along the periphery. Optimum vision is achieved when the entire eye is pointed perpendicularly towards subject matter. Minor directional adjustments appear to be achieved, without repositioning of the body, through dorsoventral and anteroposterior adjustments of the lens within the pupil (Fernald 1988).

Our study revealed that all major classes of chromatophores (melanophores, xanthophores, erythrophores, violet-blue iridophores) and crystalline platelets were present in the cornea, aqueous humor, vitreous humor, outer lens membrane and possibly the lens itself of *Poecilia reticulat*a. Contrary to visual observations and conventional transmitted light microscopy the cornea, aqueous and vitreous humor are not clear under conventional reflected light microscopy. Each possesses independent populations of static and/or free-floating chromatophores. To the authors’ knowledge this is the first time this has been reported in *P. reticulata*, though similar observations has been reported or summarized in other species either microscopically or biochemically (Dunlap 1989: Douglas 1989, 1999, 2001; Thorpe 1992; Siebeck 2001; Soules 2005; Gray 2009; Shcherbakov 2013). Both static and free-floating corneal pigments have been documented in an earlier study in humans (Snip 1981).

Ocular microscopy, subsequent to enucleation or horizontal dissection of the eye, and lens and cornea extractions, revealed that chromatophores (melanophores, xanthophores, erythrophores, and violet-blue iridophores) and also crystalline platelets were present in the tissue comprising the cornea and iris, fluids of the aqueous and vitreous humors, and the membrane surrounding the lens in both Pb and non-Pb. Collected xanthophore populations were reduced in heterozygous Pb condition and removed in homozygous condition. Clustered xanthophores, found in all parts of the body and fins in “wild-type” Pb and non-Pb, remained intact in both heterozygous and homozygous Pb condition within the eye.

Microscopic “penetration” of the cornea and past the pupil (iris and lens juncture) by ocular focusing was more difficult to achieve in non-dissected specimens, especially in homozygous Pb condition, due to a proliferation of iris melanocytes producing a “reflective sheen”. In general, this reflective sheen is noted to be more violet colored in Pb and more blue colored in non-Pb in wild-type, with much variability observed in Domestic Guppy strains vs. feral populations. The inability to penetrate the cornea by ocular focusing resulted from fully dispersed melanocytes between the cornea and lens, and / or contracted radial muscles of the iris (dilated pupils), as samples were prepared and initially viewed shortly after euthanasia.

Observations were taken 1 hour after euthanasia with both dissected and non-dissected, cornea intact and after cornea removal, allowed for greater microscopic penetration after melanocyte constriction. This revealed static melanophores in the cornea, iris and lens, and free-floating melanophores in the aqueous and vitreous humors, a dense layer of violet-blue iridophores, high xanthophore and minimal erythrophore populations. Population levels of each varied between Pb and non-Pb, among strains (populations), and within individuals. Thus, the presence of all chromatophore types should be considered the “normal” in Guppy ocular media, just as they are in the body and finnage.

The dense layer of violet-blue iridophores (Pb with a higher ratio of violet to blue, and non-Pb balanced ratio of violet to blue), under varying reflected light, was consistent over the entire pupillary region in both Pb and non-Pb. A dark violet reflective sheen over the pupil was frequently evident in Pb. In non-Pb it was often more difficult to observe this reflective sheen, and the appearance was bluer. It has long been thought that sensitivity to red and blue light are a heritable factor (Houde 1991, pg. 116, personal communication with Endler). We take this a step further to include sensitivity to UV and near-UV wavelengths as being heritable through Pb (Bias and Squire 2017a).

Results are presented in the following format and order: A. Non-dissected pupil and iris (Fig 6-8), partial dissection of eye (non-enucleated) with orbit, operculum and dentary intact viewed from high angle (Fig 9-10) and perpendicular (Fig 11), horizontal axis dissection of the eye with lower portions of orbit, operculum and dentary intact viewed from high-angle (Fig 12), B. Protruding lens intact with cornea removed (Fig 13-14), C. Corneal Extraction (Fig 15-21), D. Aqueous Humor Fluid Extraction (Fig 22-25), E. Vitreous Humor Fluid Extraction (Fig 26-28), F. Lens complete extraction (Fig 29-36). All non-dissected images were taken from the right side and all dissection was done on the left side, unless otherwise indicated. I think you need to standardize the use of upper and lower case in these sections.

**Fig 6.**
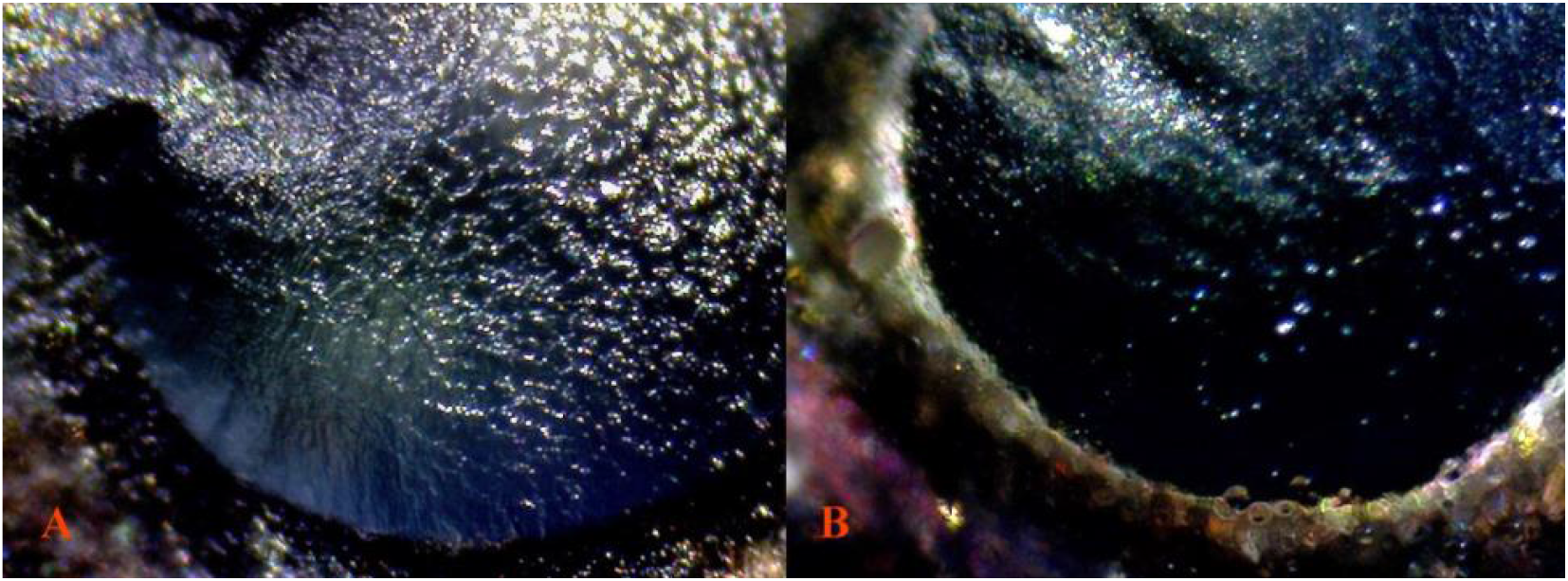
Wet mounts, no cover glass. **(A)** 18 Pb 40× 17 **Pb/Pb* (non-dissected*) reflected light. Higher violet iridophore reflective sheen in the pupillary region, producing a more “purple” appearance. Isolated xanthophore presence, either reflected or resident in pupil. **(B)** 19 non-Pb 40× 25 **Pb/Pb* (non-dissected*) reflected light. Balanced violet-blue iridophore reflective sheen in the pupillary region, producing a more “blue” appearance. Isolated single cells appear to be reflected in the pupil.

**Fig 7.**
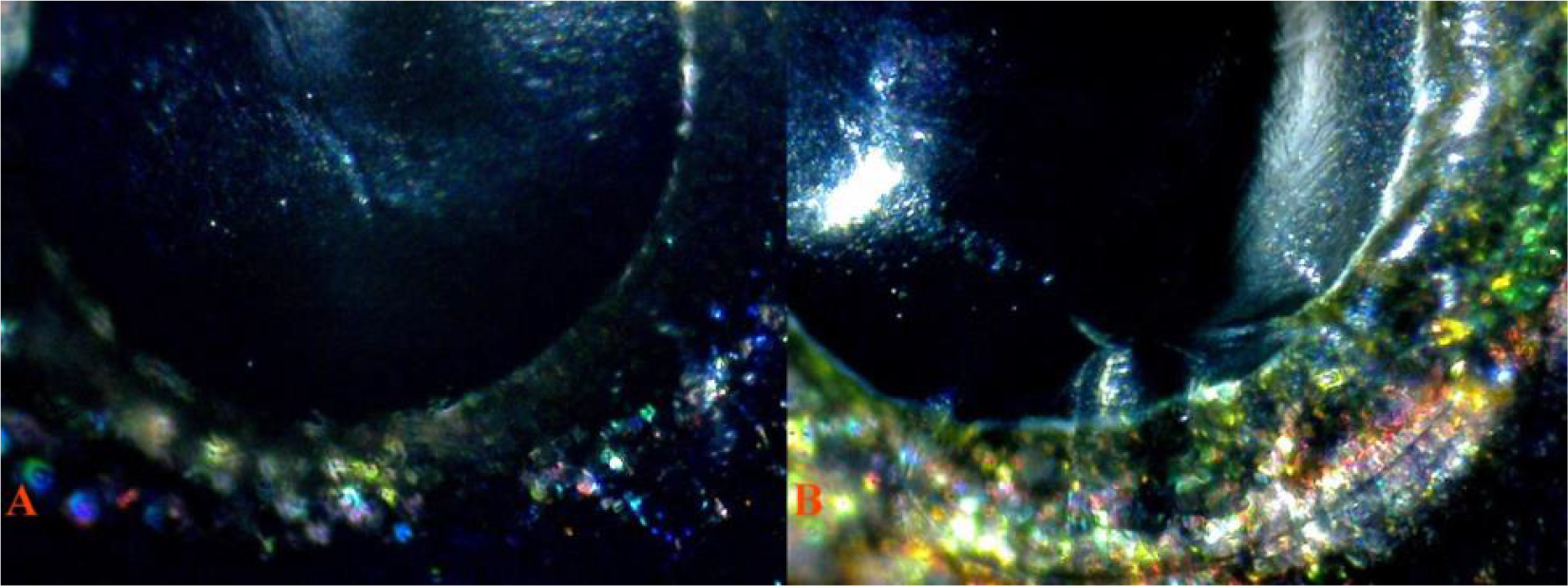
Wet mounts, no cover glass. **(A)** 23 Pb 40× 4 **Pb/Pb* (non-dissected*) reflected light. Higher violet iridophore reflective sheen in the pupillary region. Reduced xantho-erythrophore content found in the iris. **(B)** 24 non-Pb 40× 3 **Pb/Pb* (non-dissected*) reflected light. Balanced violet-blue iridophore reflective sheen in the pupillary region. Higher degree of variation in chromatophore types commonly found in non-Pb iris. Isolated single cells appear to be reflected in the pupil.

**Fig 8.**
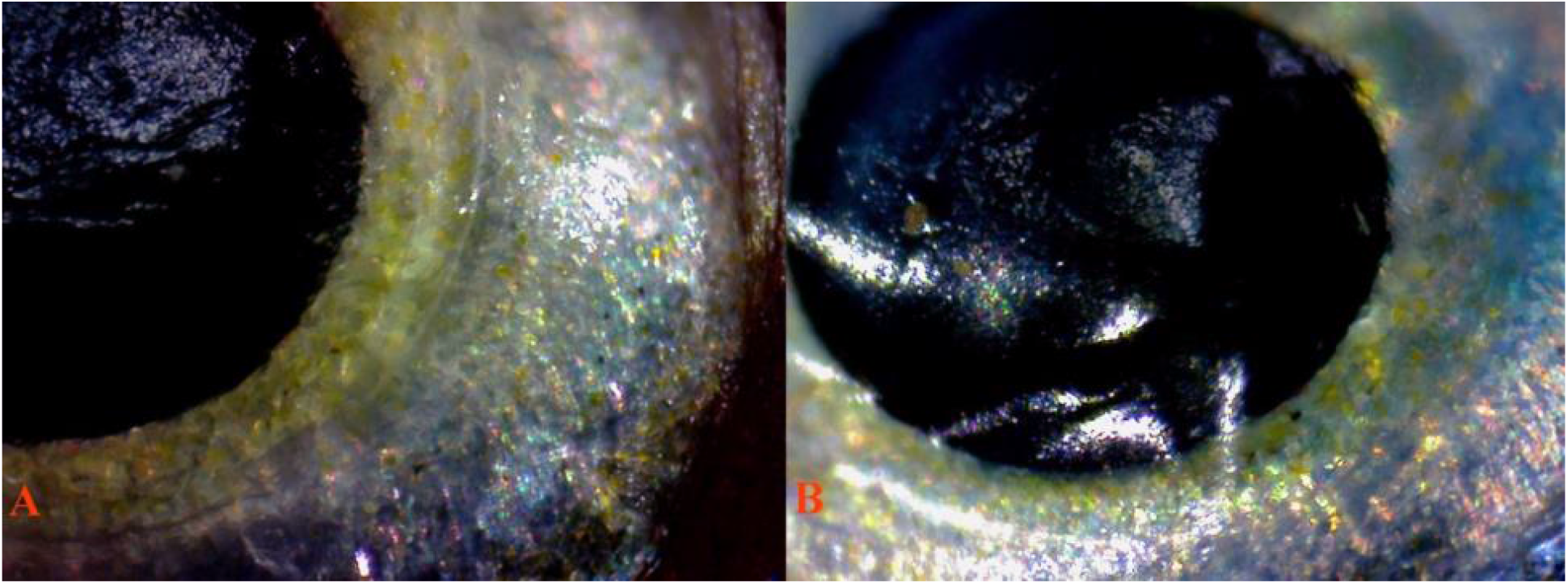
Wet mounts, no cover glass. **(A)** Blond 29 Pb 40× 21 **Pb/Pb* (non-dissected*) reflected light. Higher violet iridophore reflective sheen in the pupillary region, producing a more “purple” appearance. Reduced melanophore size (caused by the Blond mutation) visible in iris and likely similarly in the pupil region. **(B)** Blond 28 non-Pb 40× 13 **Pb/Pb* (non-dissected*) reflected light. Balanced violet-blue iridophore reflective sheen in the pupillary region, producing a more “blue” appearance. Reduced melanophore size visible in iris and likely similar in the pupil region. Isolated single cells appear to be reflected in the pupil.

**Fig 9.**
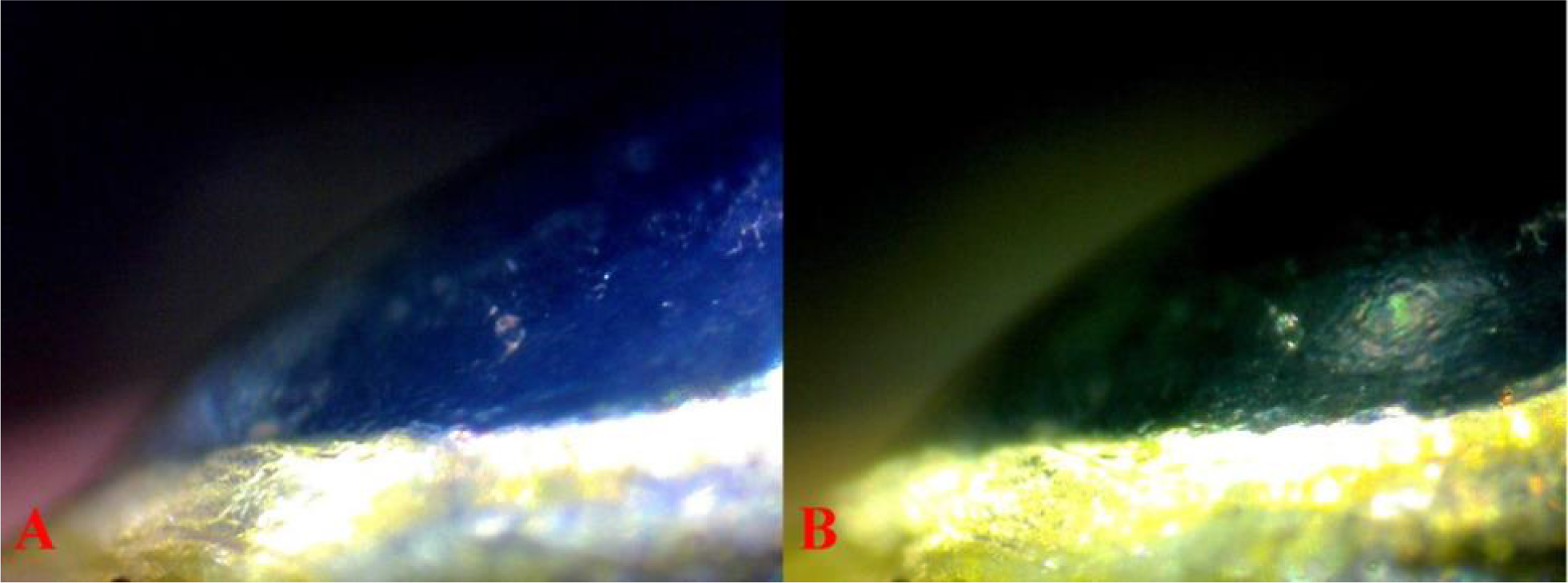
Wet mounts, no cover glass. **(A)** 31 40× 9 **Pb/Pb* (partial dissection*) low angle reflected light revealing iridophore reflective sheen in the pupillary region. **(B)** The same field, high angle reflected light revealing xanthophore reflective sheen in the pupillary region.

**Fig 10.**
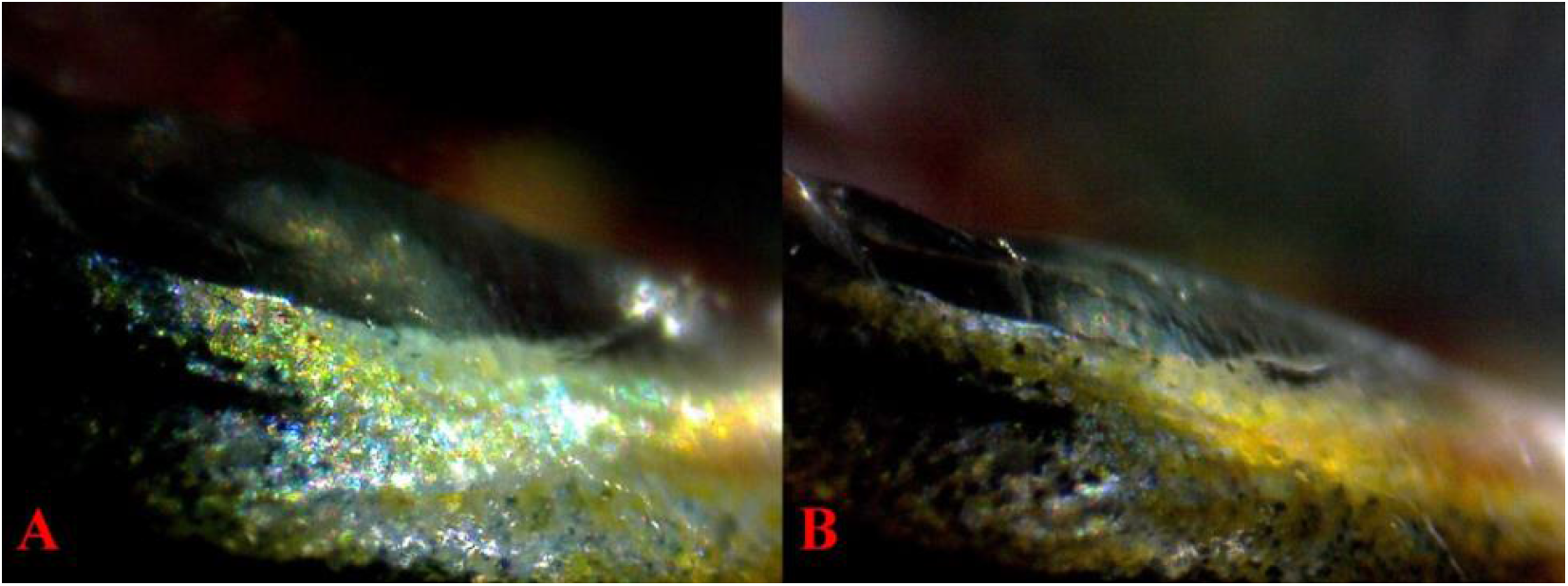
Wet mounts, no cover glass. **(A)** 31 40× 11 **Pb/Pb* (partial dissection*) reflected light. **(B)** 31 40× 19 **Pb/Pb* (partial dissection*) reflected light. Both images clearly show xanthophore reflection from the cornea into the pupillary region.

**Fig 11.**
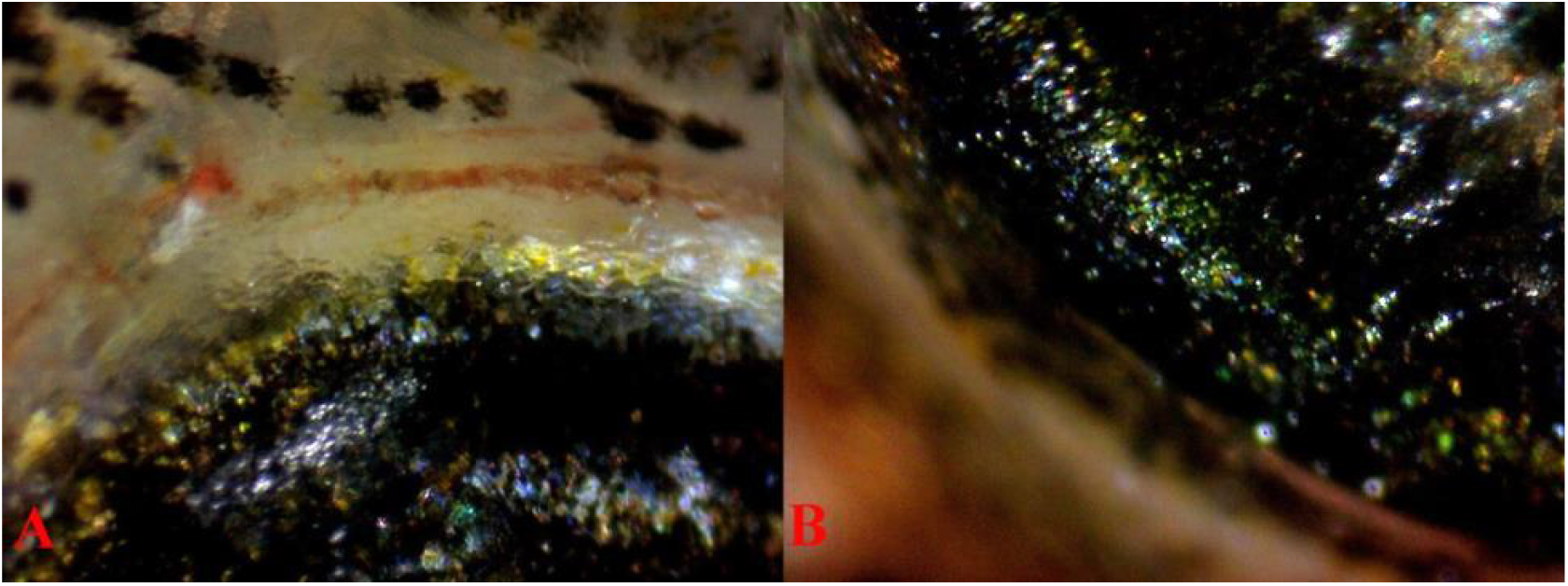
Wet mounts, no cover glass. **(A)** 30 40× 3 **Pb/Pb* (partial dissection*) reflected light, revealing chromatophore populations in the iris and scleral ring. **(B)** 30 40× 5 **Pb/Pb* (partial dissection*) reflected light revealing chromatophore population in the iris. Each photo appears to show actual pigment cells in the cornea above the pupil.

**Fig 12.**
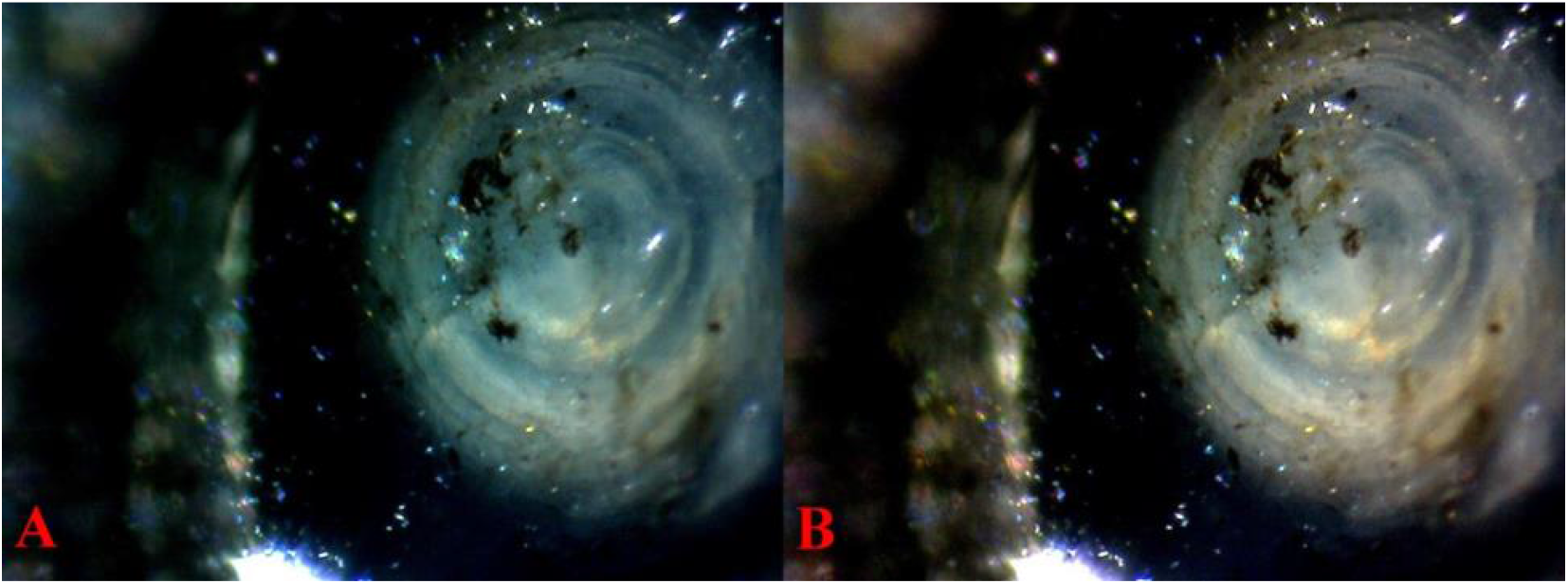
Wet mounts, no cover glass. **(A)** 32 40× 5 *Pb/- (partial dissection*) reflected light, high angle. **(B)** The same field, reflected light with white balance adjusted. Superficial free-floating chromatophores are visible in the aqueous-vitreous humor. The cells over the lens appear to have been dislodged from vitreous humor. The “dark-matter” over the dissected lens plane appears similar, rather than inclusions within the actual lens. Reflection through the pupil opening is visible in lower left portion of images, from which the lens has retracted. Approximately 60% of the ventral portion of the eye remains within the orbit.

**Fig 13.**
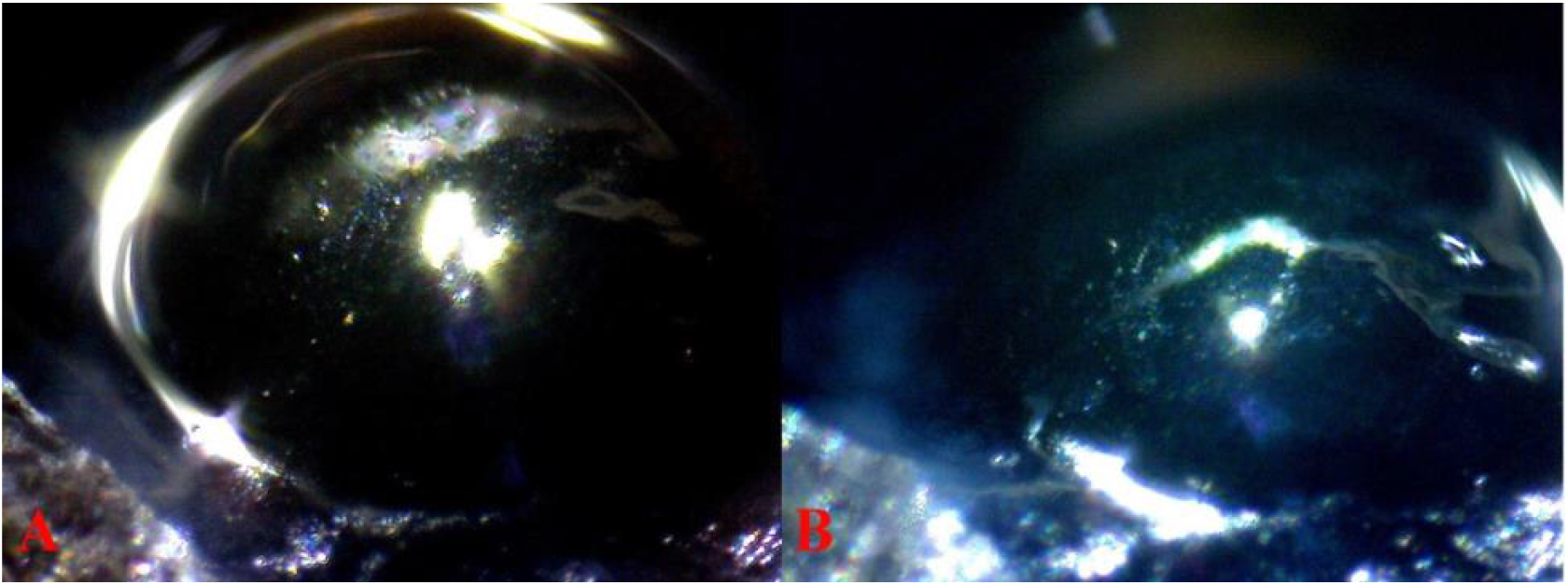
Wet mounts, no cover glass. **(A)** 32 40× 15 *Pb/- (partial dissection*) low angle reflected light with white balance adjusted. Image taken at 45 degree angle. **(B)** The same field, reflected light with no white balance adjustment. In both images a small portion of the surrounding capsule of the lens anterior pole is partially missing after scleral skin and true cornea removal. The missing section can be seen in lower left of **Fig 17A** and **Fig 18A.** A xanthophore and violet-blue iridophore reflected sheen is seen in the protruding lens, over regions with and without the surrounding capsule.

**Fig 14.**
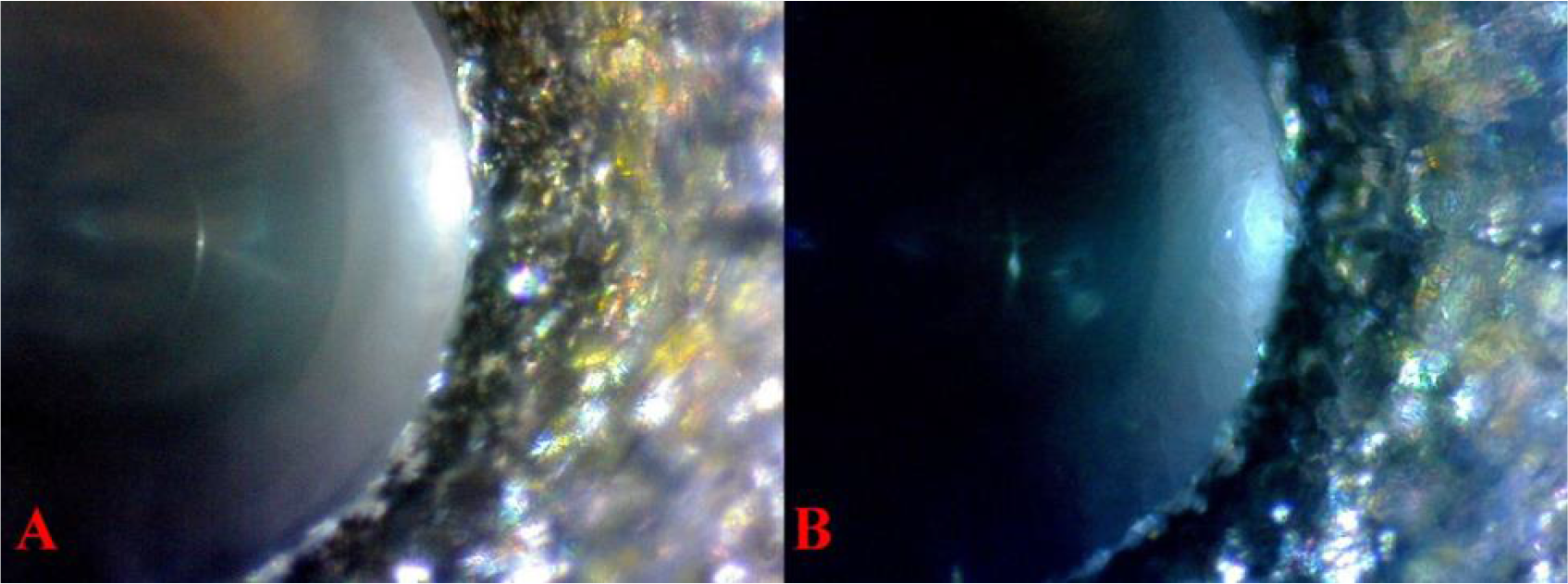
Wet mounts, no cover glass. **(A)** 32 40× 10 *Pb/- (partial dissection*) high angle reflected light with white balance adjusted. **(B)** The same field, reflected light with no white balance adjustment. Lacking a cornea, xanthophore and violet-blue iridophore reflective sheen is “muted” as seen over the the protruding lens, and no reflected single cells are visibile. In each view the lens nucleus is visible.

**Fig 15.**
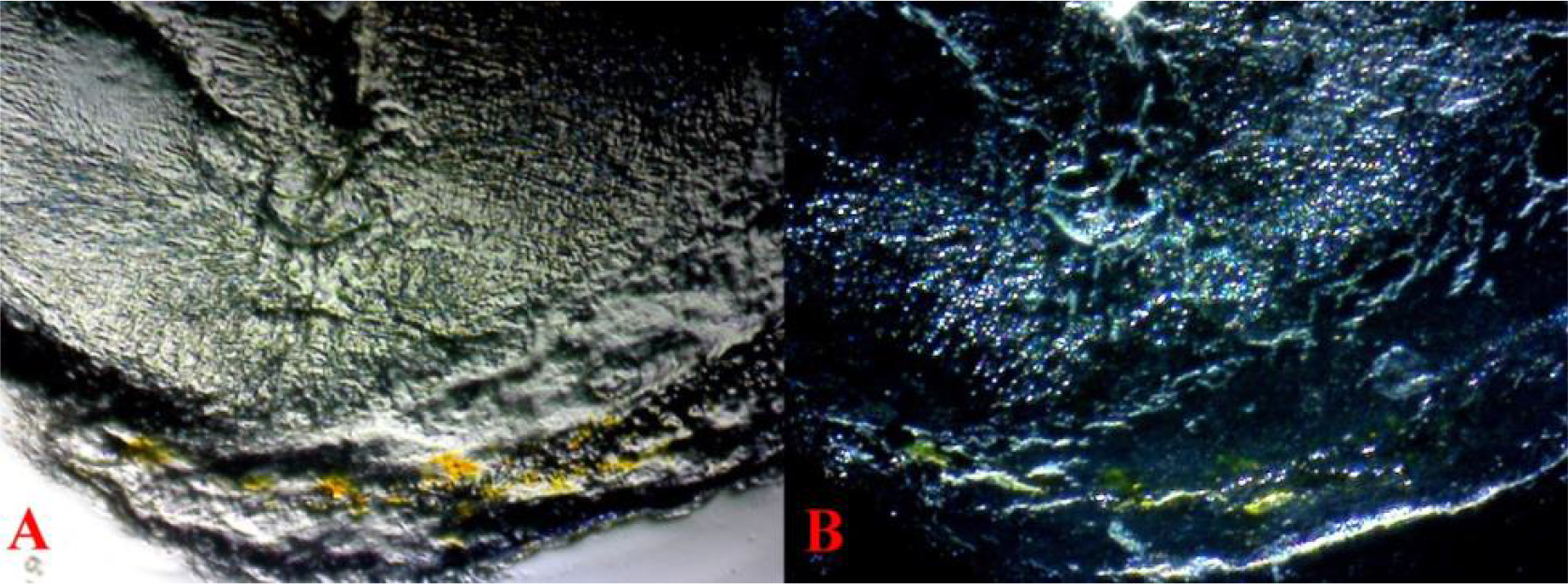
Wet mounts, no cover glass. Corneal extraction **(A)** 32 40× 2 *Pb/- (dissection*) reflected and transmitted light with white balance adjusted. **(B)** The same field, reflected light with no white balance adjustment.

**Fig 16.**
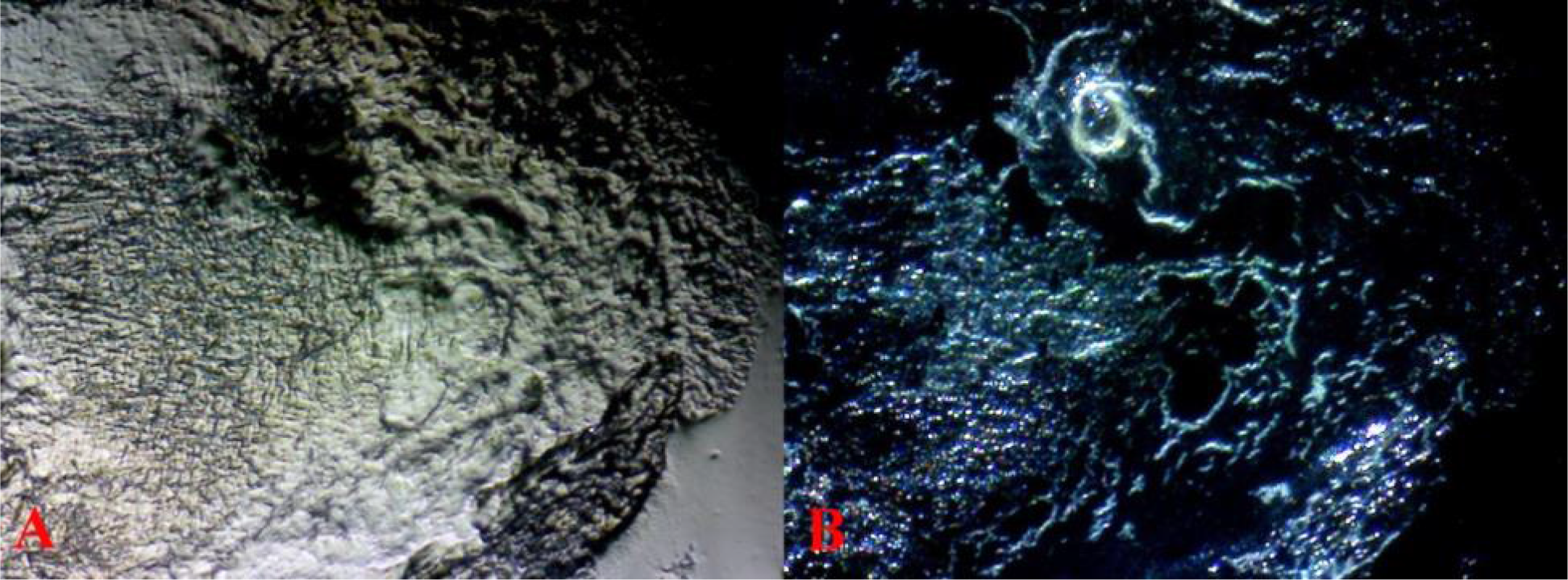
Wet mounts, no cover glass. Corneal extraction **(A)** 32 40× 4 *Pb/- (dissection*) reflected and transmitted light with white balance adjusted. **(B)** The same field, reflected light with no white balance adjustment.

**Fig 17.**
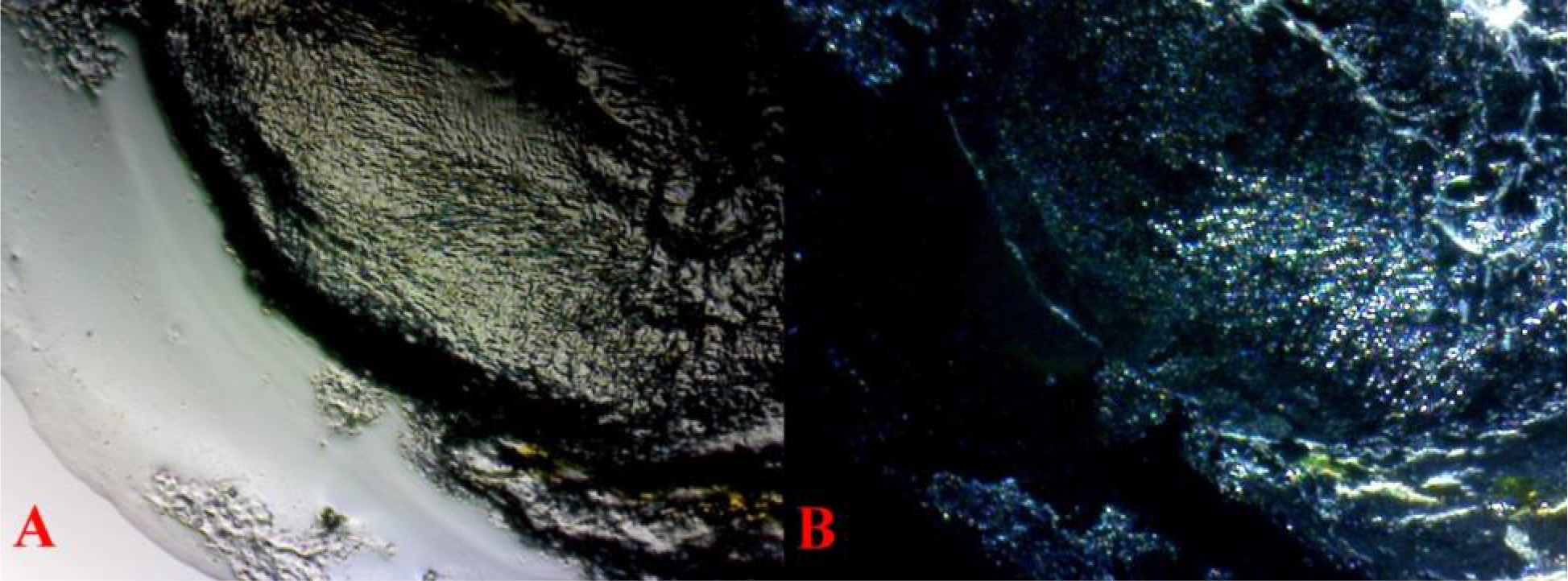
Wet mounts, no cover glass. Corneal extraction **(A)** 32 40× 6 *Pb/- (dissection*) transmitted light with white balance adjusted. **(B)** The same field, reflected light with no white balance adjustment. Single cells within exterior clear tissue (lower left) are free-floating from the aqueous humor, larger clusters are attached iris tissue (lower center), along with large area of xanthophores. When hydrated the clarity of true cornea is less visible in the central portion of the image with increased visibility of the attached underlying portion of the surrounding capsule of the lens. Violet-blue iridophores are visible in the central portion of the image (underlying the scleral and true cornea) under transmitted light and under reflected light. This indicates both endothelial positioning and free-floating cells. Black areas under reflected light are clear cornea devoid of any attached underlying tissue.

**Fig 18.**
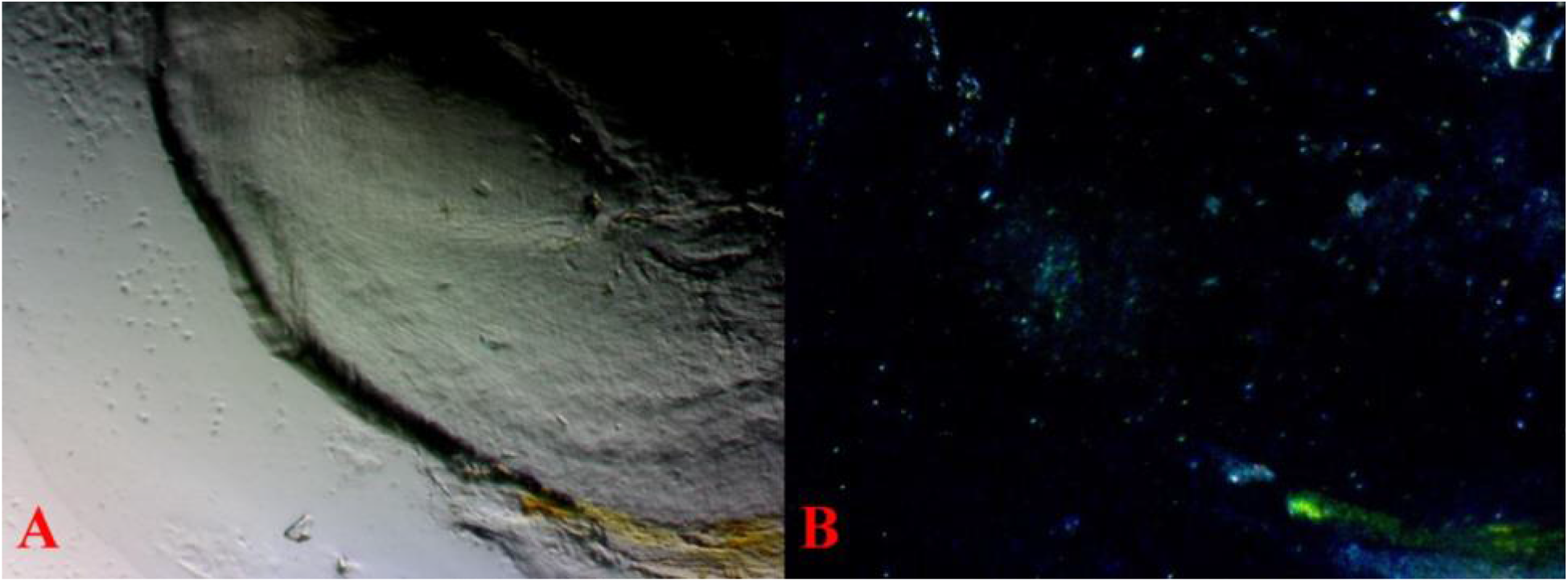
Prepared mounts with cover glass. Corneal extraction **(A)** 32 40× 8 *Pb/-(dissection*) transmitted light with white balance adjusted. **(B)** The same field, reflected light with no white balance adjustment. Single cells within exterior clear tissue (lower left) are free-floating from the aqueous humor, larger clusters are attached iris tissue (lower center), along with a large area of xanthophores. When prepared as a permanent mount the clarity, the true cornea is visible in the central portion of the image with decreased visibility of the attached underlying portion of the surrounding capsule of the lens (thick white area). No reflective violet-blue iridophores are visible in the central portion of the image (underlying scleral and true cornea) under transmitted light, and visibly reduced under reflected light. This indicates both endothelial positioning and free-floating cells.

**Fig 19.**
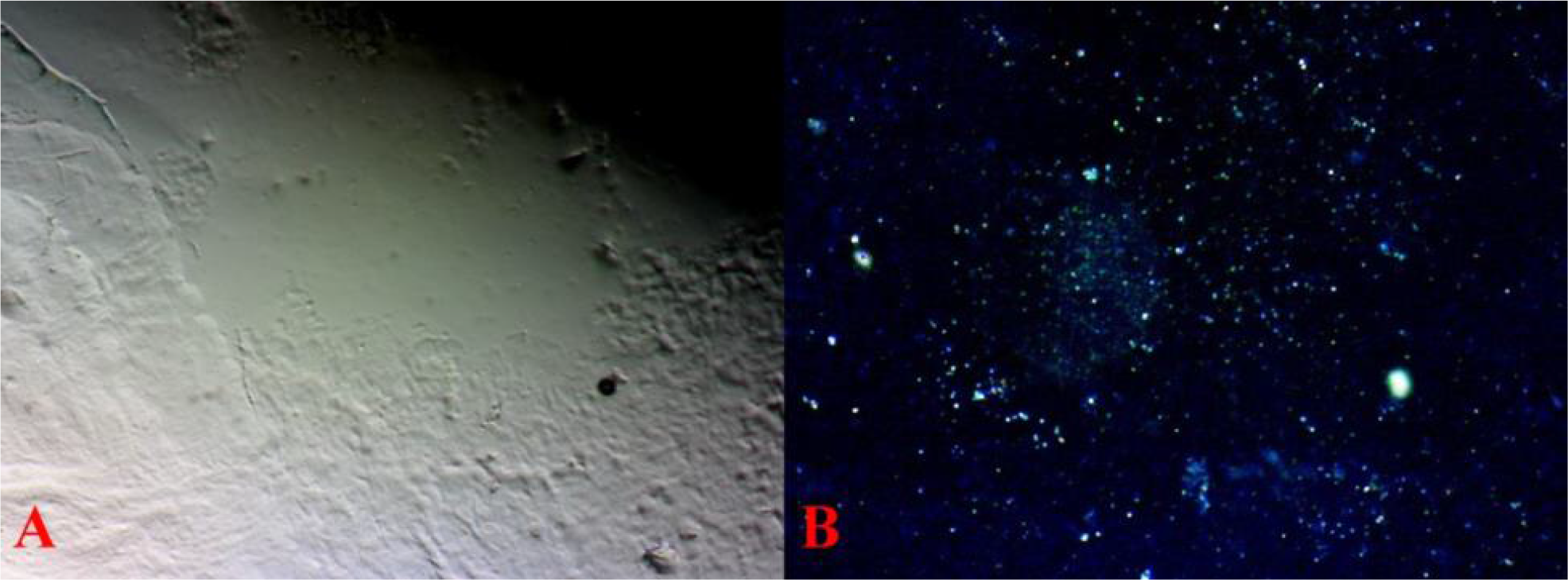
Prepared mounts, with cover glass. Corneal extraction **(A)** 32 100× 9 *Pb/-(dissection*) transmitted light with white balance adjusted over central portion of the cornea. **(B)** The same field, reflected light with no white balance adjustment. When prepared as permanent mount at higher resolution the clarity of the true cornea is not visible in the image, only the visibly attached underlying portion of the surrounding capsule of the lens (thick white). No reflective violet-blue iridophores are visible under transmitted light, and visibly reduced under reflected light. This indicates both endothelial positioning and free-floating cells, as there is clear distinction in size of the latter being at a high level.

**Fig 20.**
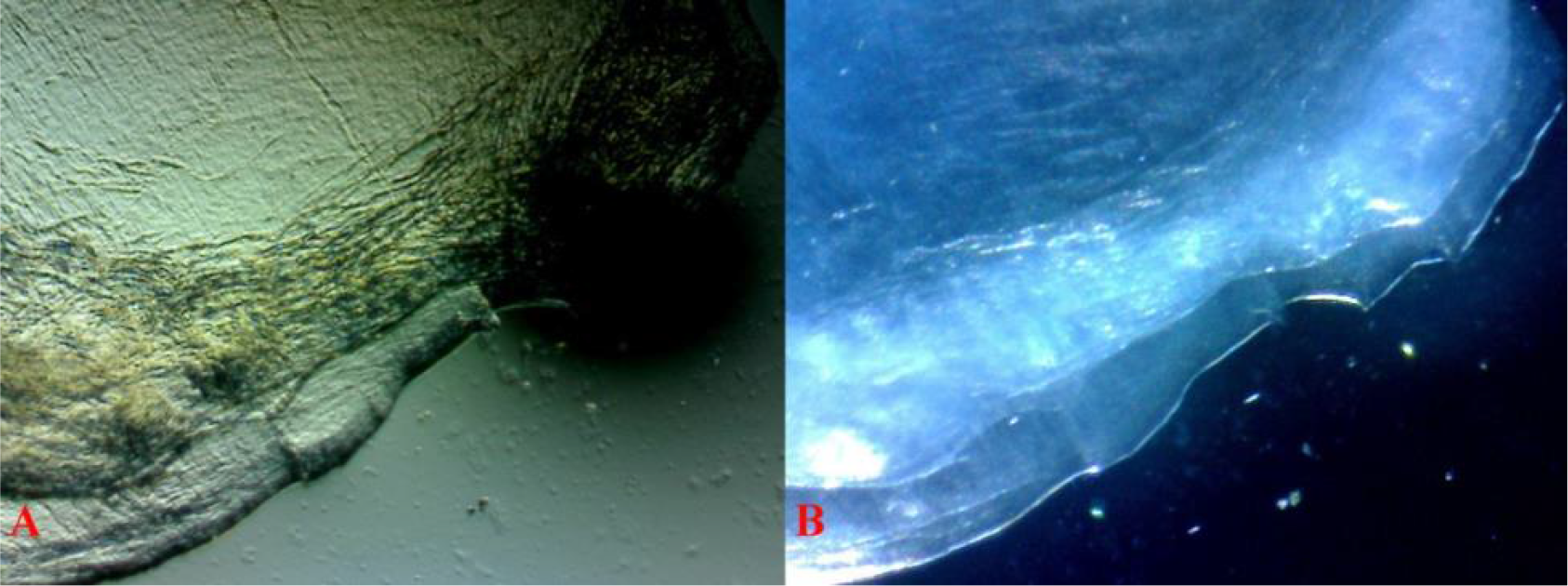
Wet mounts, no cover glass. Corneal extraction **(A)** 34 40× 3 *pb/- (dissection*) transmitted light over the central portion of the cornea. **(B)** The same field, reflected light. Thin dissection over the central pupillary region, lacking surrounding scleral or iris tissue. While the appearance is much clearer, minimal endothelial chromatophores are suspended with minimal numbers of free-floating cells visible on the slide beyond the corneal tissue under both transmitted and reflected light.

**Fig 21.**
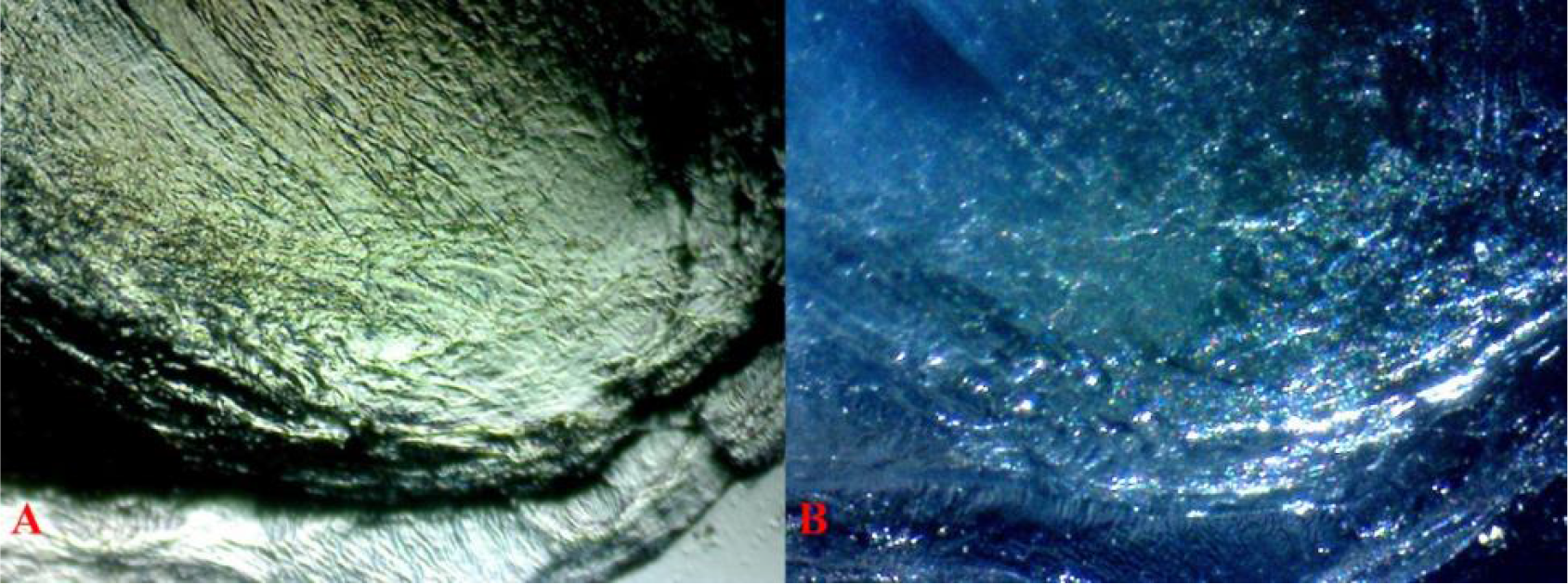
Wet mounts, no cover glass. Corneal extraction **(A)** 34 40× 10 *pb/- (dissection*) transmitted light over the central portion of the cornea. **(B)** The same field, reflected light. Thin dissection over the central pupillary region, lacking surrounding scleral or iris tissue. While appearance is much clearer minimal endothelial chromatophores are suspended with higher numbers of free-floating cells visible on the slide beyond the corneal tissue under both transmitted and reduced reflected light.

**Fig 22.**
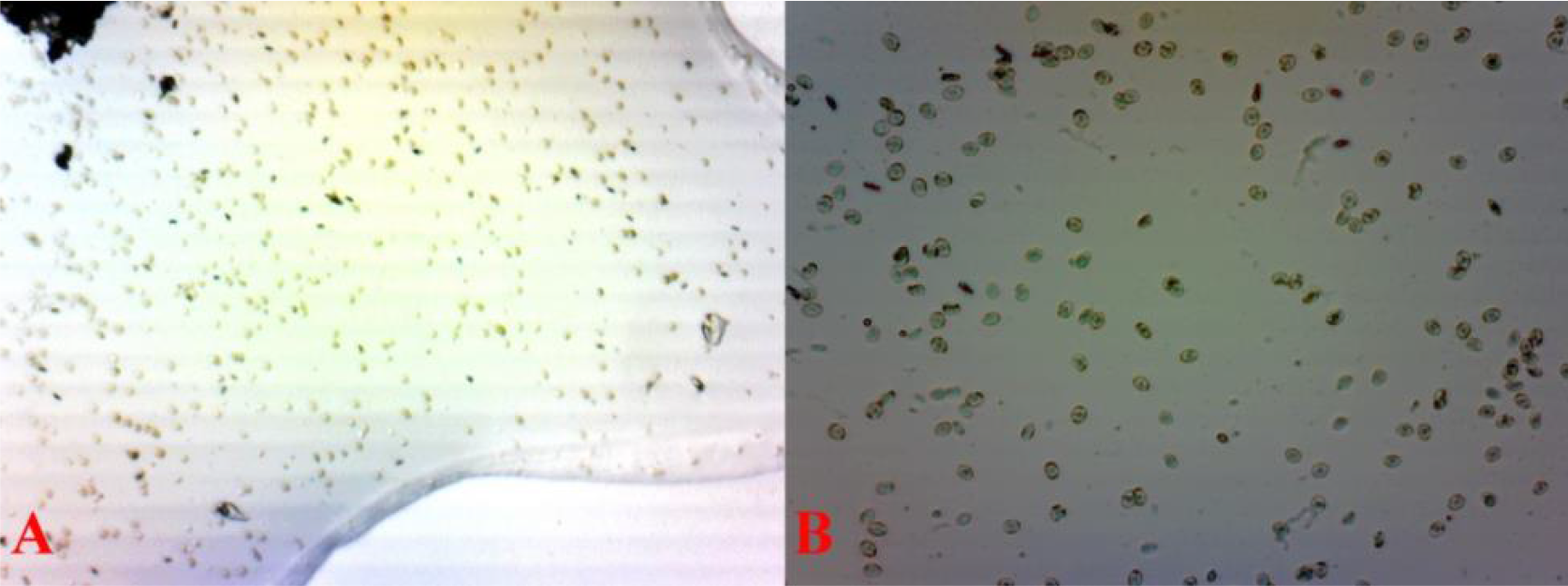
Wet mounts, no cover glass. Aqueous humor fluid extraction **(A)** 32 40 6 *Pb/-(dissection*) reflected and transmitted light. **(B)** 32 100 7 *Pb/- (dissection*) transmitted light with white balance adjusted.

**Fig 23.**
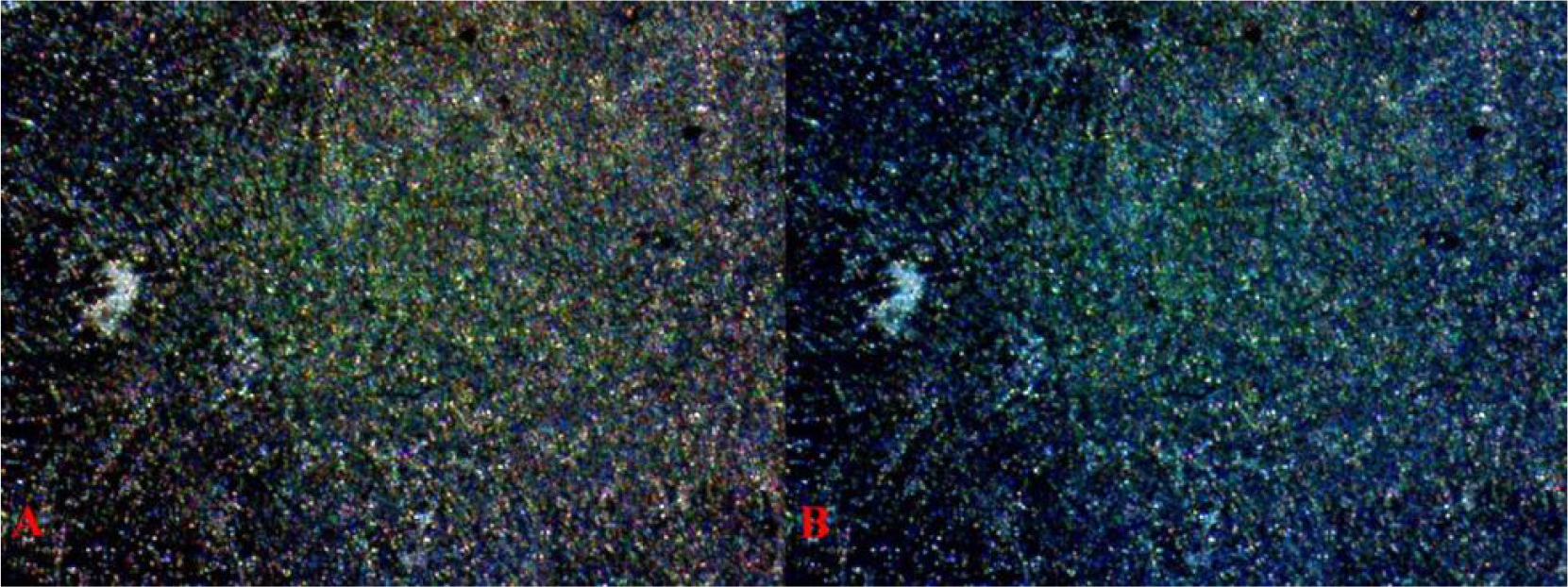
Wet mounts, no cover glass. Metal Gold (Mg) male. Dehydrated aqueous humor fluid extraction **(A)** 36 40 2 **Pb/Pb* (dissection*) reflected light with white balance adjusted. Notice increased xanthophore population. **(B)** The same field reflected light. Notice more blue appearance with balanced violet-blue iridophores.

**Fig 24.**
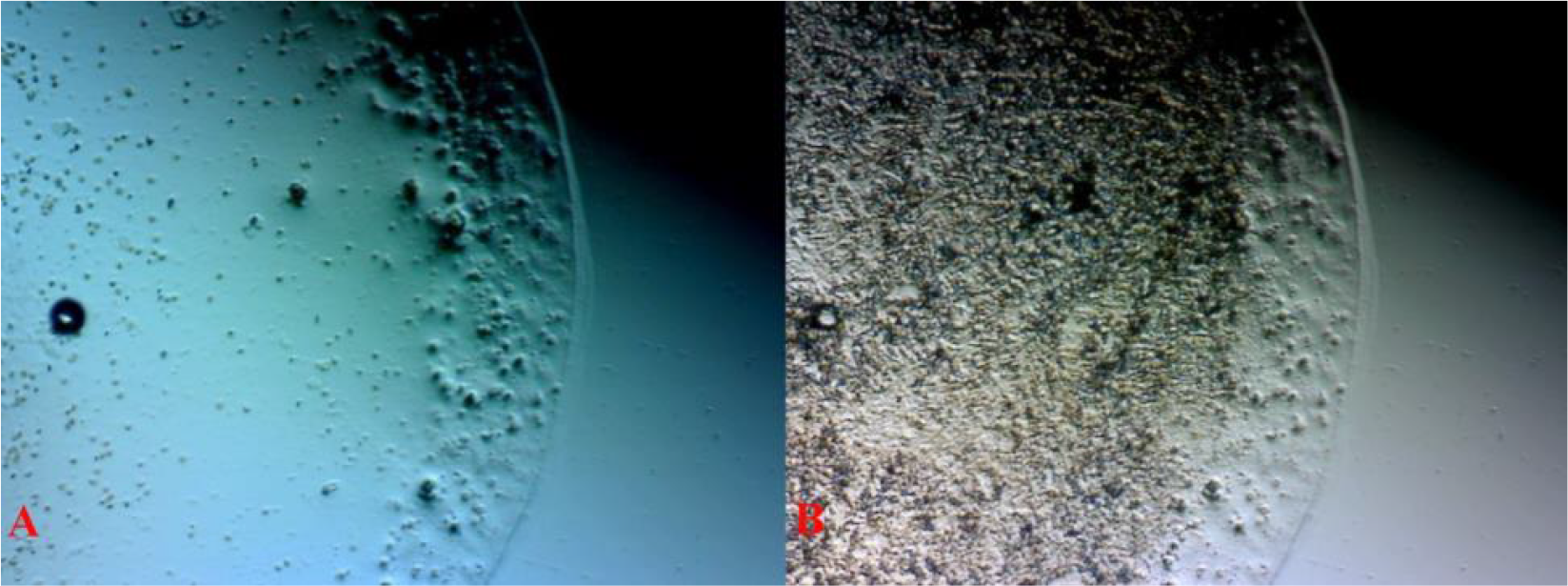
Wet mounts, no cover glass. Dehydrated aqueous humor fluid extraction **(A)** 33 40 1 *Pb/- (dissection*) transmitted light. **(B)** The same field transmitted light with white balance adjusted.

**Fig 25.**
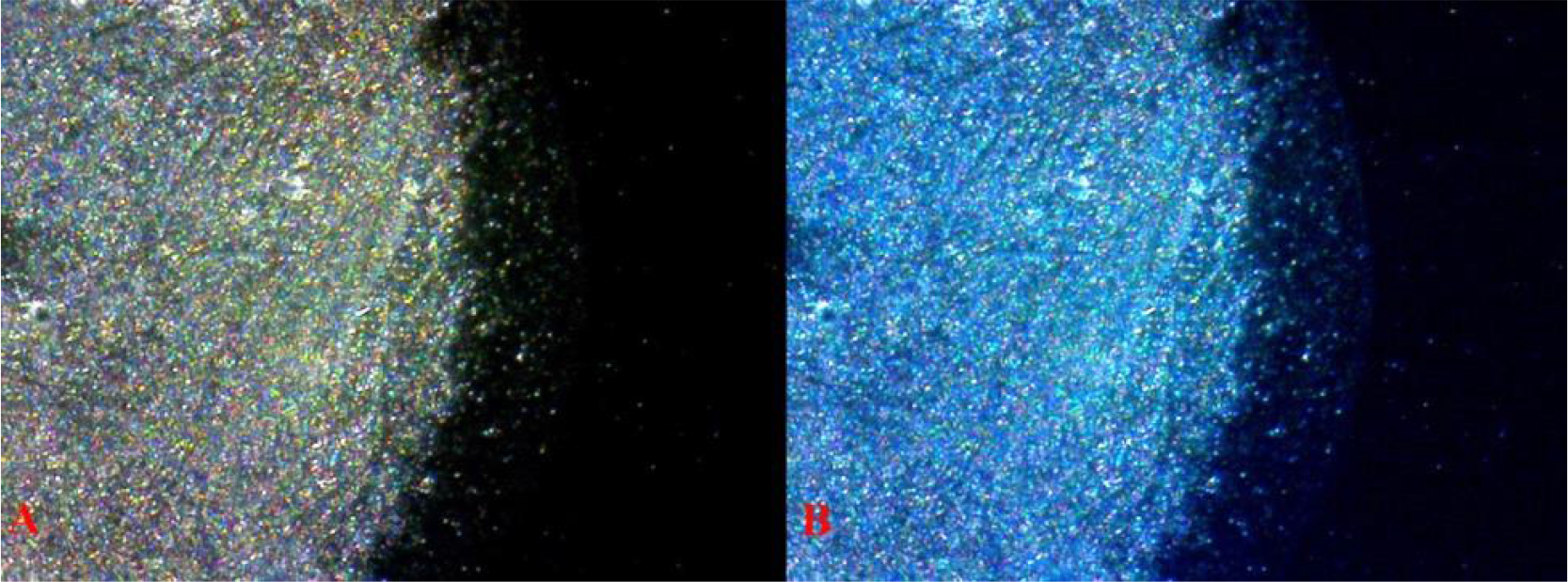
Wet mounts, no cover glass. Dehydrated aqueous humor fluid extraction **(A)** 33 40 1 *Pb/- (dissection*) reflected light with white balance adjusted. **(B)** The same field reflected light.

**Fig 26.**
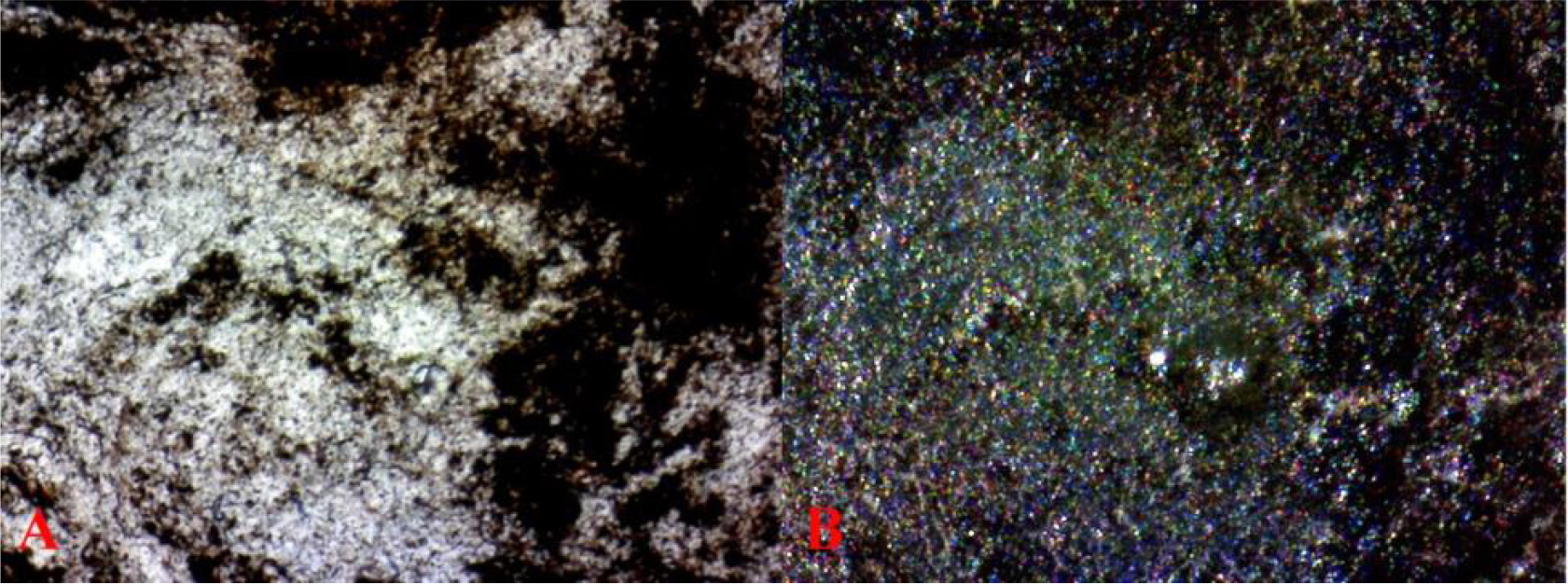
Wet mounts, no cover glass. Vitreous humor dehydrated **(A)** 31 40× 1 **Pb/Pb** transmitted light with white balance adjusted. **(B)** The same field, reflected light with white balance adjusted. Notice xanthophores are present with more violet expression form higher concentration of violet iridophores

**Fig 27.**
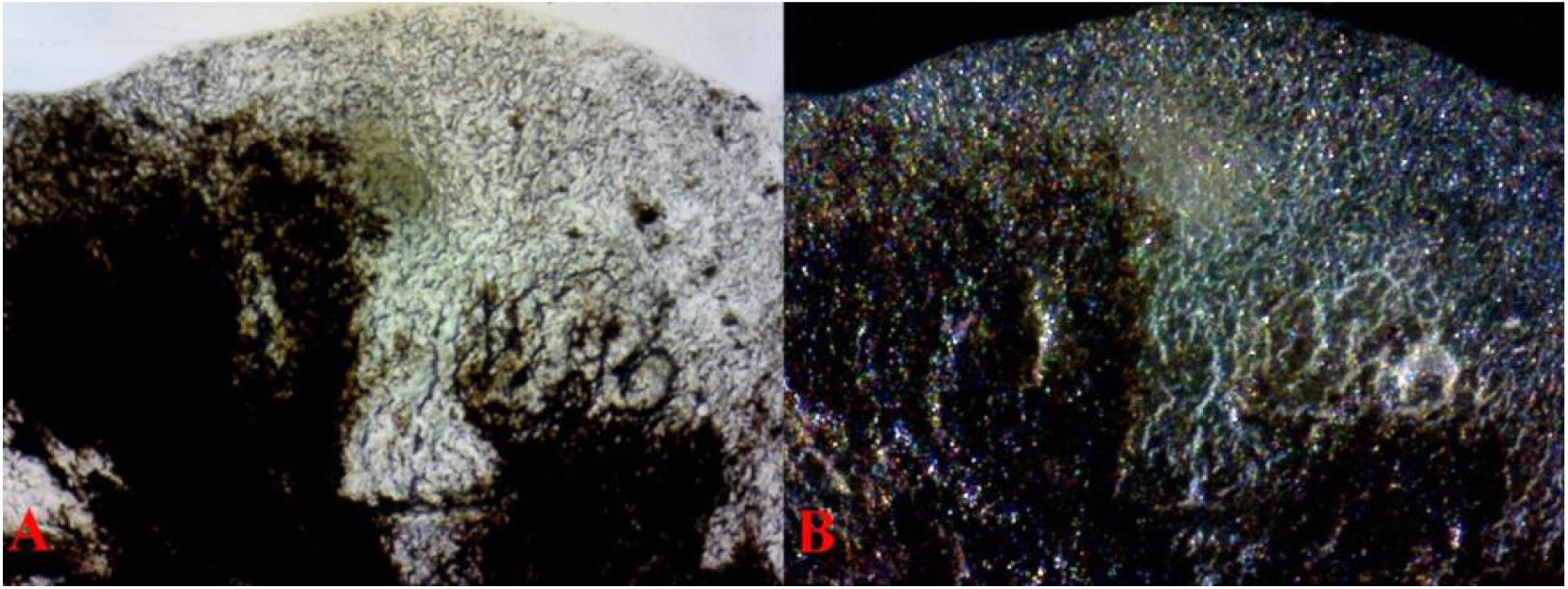
Wet mounts, no cover glass. Vitreous humor partially dehydrated **(A)** 31 40× 3 **Pb/Pb** transmitted light with white balance adjusted. **(B)** The same field, reflected light with white balance adjusted. Notice xanthophores are present with more violet expression form higher concentration of violet iridophores

**Fig 28.**
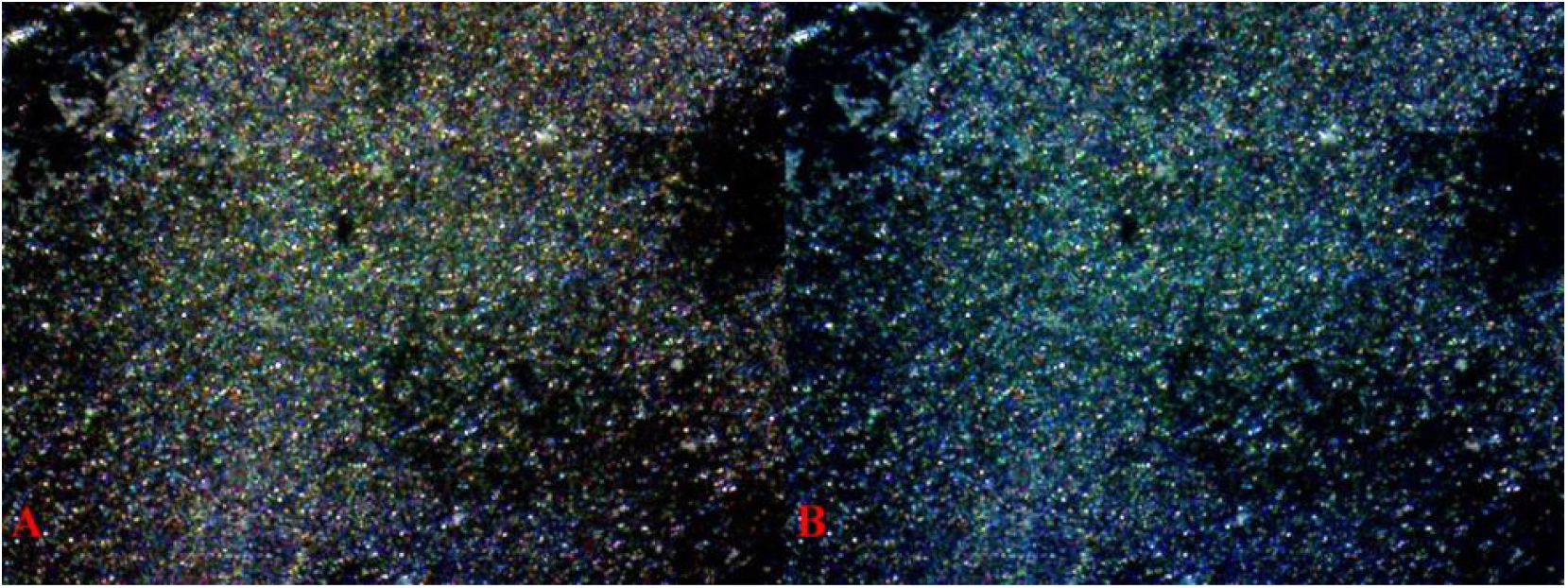
Wet mounts, no cover glass. Metal Gold (*Mg*) male. Dehydrated aqueous humor fluid extraction **(A)** 36 40 2 **Pb/Pb* (dissection*) reflected light with white balance adjusted. Notice the increased xanthophore population corresponding to the Mg mutation. **(B)** The same field reflected light. Notice more blue appearance with balanced violet-blue iridophores. Both photos express higher melanophore populations as compared to prior aqueous humor images.

**Fig 29.**
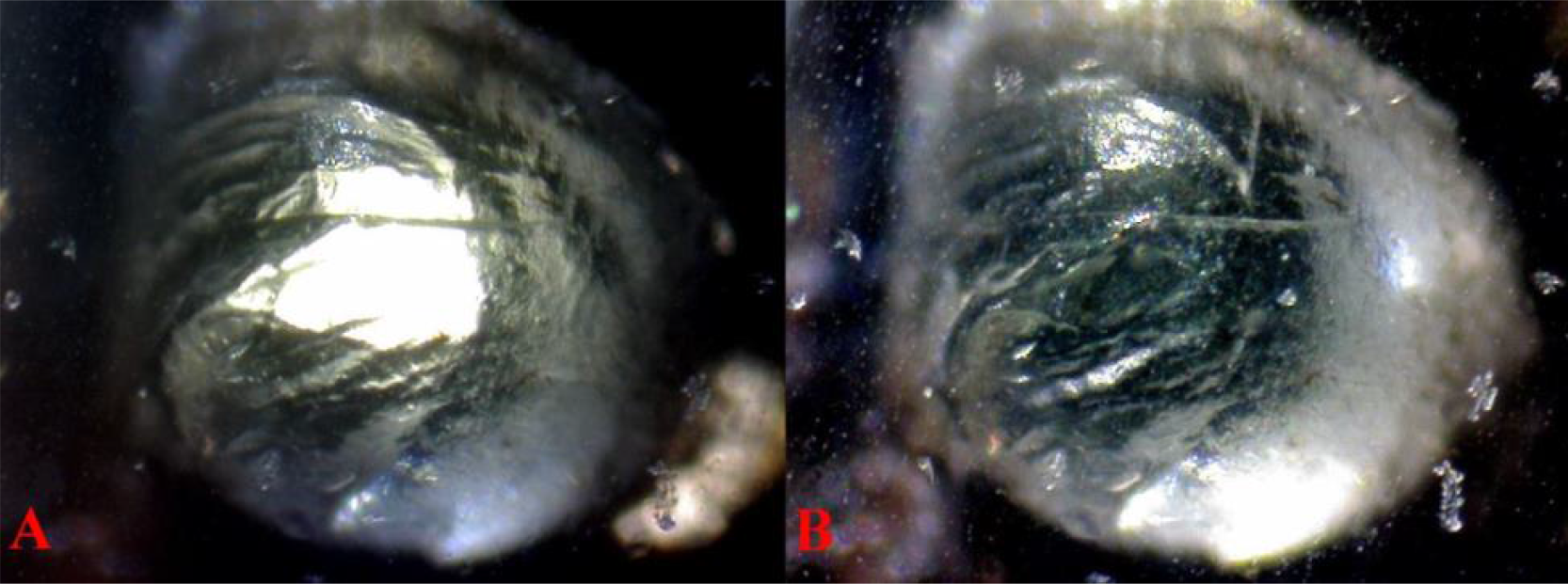
**(A)** 17 40× 7 **Pb/Pb* (full dissection and partial extraction*) reflected and transmitted light. **(B)** The same field, transmitted light.

#### A. Cellular Comparison: Iris and Pupil pigmentation

***I***. Non-dissected samples (Fig 6–8) used full body specimens to incorporate complete spectral qualities of all chromatophore populations, both within and surrounding the eye. All images were under reflected lighting. ***II***. Partial dissections (Fig 9–11) were on the left lateral side with incision along the median plane of the skull from mouth to supraocciptal, followed by dorsoventral removal to include the complete operculum and eye intact within the bony orbit. All images were under low angle reflected lighting. ***III***. Horizontal axis dissection (Fig 12) of the eye (frozen) approximately mid-level with lower portions of orbit, operculum and dentary intact viewed from high-angle under reflected lighting. A resident population of chromatophores in the iris and their reflection into the pupillary region confirms the presence of an ocular media filter mechanism in *P. reticulata*.

#### B. Cellular Comparison: Protruding Lens Intact With True Cornea Removed

The removal of a portion of the scleral skin and the entire true cornea was performed on frozen specimens (Fig 13–14) to preserve the integrity of the structure and maximize the amount of cornea removed. Freezing of ocular media has been shown to have no significant effect (Douglas 2014). Extraction was performed from the right side of body with the incision made anterior to the clear central area of the pupil and extending slightly into the posterior side iris, providing complete exposure of the protruding lens. A resident population of chromatophores in the iris after corneal removal and their reflection into the pupil region again confirms the presence of an ocular media filter mechanism in *P. reticulata*.

#### C. Cellular Comparison: Corneal Extraction

The removal of a portion of scleral skin and entire true cornea was performed on frozen specimens to preserve the integrity of the structure and maximize the amount of cornea removed. Freezing of ocular media has been shown to have no significant effect (Douglas 2014). Extraction was performed from the right side of body with an incision made anterior to the clear central area of the pupil and extending slightly into the posterior side of the iris. An effort was made to remove a very thin section of upper most exterior clear tissue and continuing into deep tissue of the true cornea to include all its layers (epithelium to endothelium), by cutting horizontally across the natural convex curvature of the eye. As desired, some iris tissue appears to be present on the exterior ventral portion of the samples. After removal the cornea was rinsed multiple times in saline solution, with prolonged soaking, to remove possible free-floating chromatophore contamination. When the cornea was viewed under a high magnification hand lens under transmitted light, no discoloration was apparent over the entire sample; i.e. it appeared clear. Microscopic results show otherwise.

In images (Fig 15, 17-18) on the lower exteriors is a thin section of clear tissue, in white **(A)** and black **(B),** from over the scleral layer of the cornea that extends over the entire underlying true cornea. The circular outline of cornea is visible to varying degrees. When enlarged, violet-blue iridophores and xantho-erythrophores are seen in the central portion of the cornea, which is comprised of all layers (epithelium to endothelium). Cells are believed to primarily reside in endothelial tissue and / or are attached to anterior portions of the surrounding capsule of the lens. Also minimally present are free-floating chromatophores of the aqueous humor not removed during multiple rinsings. Xantho-erythrophores, melanophores and iridophores in the central lower portion of the image are contained in attached underlying iris tissue. In image (Fig 19) at higher magnification and prepared as a permanent mount, the clarity of the true cornea is not visible, only the attached underlying portion of surrounding capsule of the lens (thick white). No reflective violet-blue iridophores are visible under transmitted light, and are visibly reduced in number under reflected light. Some cell positions can be identified under transmitted light, this suggests the majority are endothelial vs. epithelial positioned. Images (Fig 20-21) are of a single-rinsed thin dissection over the central pupillary region, lacking surrounding scleral or iris tissue. A single rinse was used to potentially distinguish between resident endothelial cells and free-floating cells above the cornea, below the cornea and beyond the cornea over the empty area of slide. While the appearance is much clearer, minimal endothelial chromatophores are suspended with higher numbers of free-floating cells visible on slide beyond the corneal tissue under both transmitted and reduced reflected light.

#### D. Cellular Comparison: Aqueous Humor Fluid Extraction

Multiple samples of aqueous humor fluid, from several specimens, were obtained by low angle extraction with a 1ml BD Micro-Fine Insulin syringe. Each sample was compared against the others for vitreous contamination. The clearest of these were utilized in the microscopy study (Fig 22–25). The aqueous humor is generally clear under transmitted light with visible free-floating cells present. Consistency is slightly gelatinous from collagen and hyaluronic acid content. Under reflected lighting the true composition of the fluid is revealed in the form of a mind-boggling proliferation of violet-blue iridophores and xantho-erythrophores. Minimal amounts of dark matter were observed in some samples. It is possible they were pulled through the iris-lens juncture during the extraction process. A resident population of chromatophores in the aqueous humor fluid again confirms the presence of an ocular media filter mechanism in *P. reticulata*. The transmitted light views provide a rough estimate in terms of how much light is able to actually pass through the chromatophore smear, while the reflected light views of the same microscopic field provide an appreciation for the density and variety of chromatophores in the aqueous humor. The light that passes through the chromatophore smear provides a rough estimate of the amount of light that could also pass through the aqueous humor on its way to the retina. It thus becomes apparent that while the chromatophore population is quite dense, it does not prevent considerable light from reaching the retina.

#### E. Cellular Comparison: Vitreous Humor Fluid Extraction

Multiple samples of vitreous humor fluid, from several specimens, were obtained by deep penetration extraction with a 1ml BD Micro-Fine Insulin syringe. Each was compared against the other for consistency of type, and several selected for microscopy study (Fig 26-28). Dissimilar to the aqueous humor, the vitreous humor is generally a mix clear fluid and large amounts of dark matter under transmitted and reflected light with visible free-floating cells present. Under reflected lighting the true composition of the fluid is revealed in the form of a mind-boggling proliferation of violet-blue iridophores and xantho-erythrophores. A resident population of chromatophores in the vitreous humor fluid again confirms the presence of an ocular media filter mechanism in *P. reticulata*. Again, the transmitted light views provide a rough estimate in terms of how much light is able to actually pass through the chromatophore smear, while the reflected light views of the same microscopic field provide an appreciation for the density and variety of chromatophores in the vitreous humor. The light that passes through the chromatophore smear provides a rough estimate of the amount of light that could also pass through the vitreous humor on its way to the retina. It thus becomes apparent that while the chromatophore population is quite dense, it does not prevent considerable light from reaching the retina.

#### F. Cellular Comparison: Lens Extraction

Micro-dissection of the lens was performed on several enucleated eyes (Fig 29-36). After complete lens extraction each lens was saline rinsed multiple times in ethyl alcohol with prolonged soaking to remove potential loose surface contaminants adhering during extraction. Microscopy was performed with the surrounding membrane intact, partially ruptured, and removed. The lens was revealed to be semi-transparent and highly reflective under reflected light. While “chromatophore color” was not reliably detected within the crystalline lens, it was consistently observed in tissue or fragments of the surrounding membrane adhering to the anterior pole of the lens. Melanophores, violet-blue iridophores and xanthophores were seen in these tissue fragments. Chromatophore presence in the surrounding membrane indicates chromatophore populations residing internal to the aqueous humor fluid and iris. A resident population of chromatophores in the surrounding membrane of the lens again confirms the presence of an ocular media filter mechanism in *P. reticulata*.

**Fig 30.**
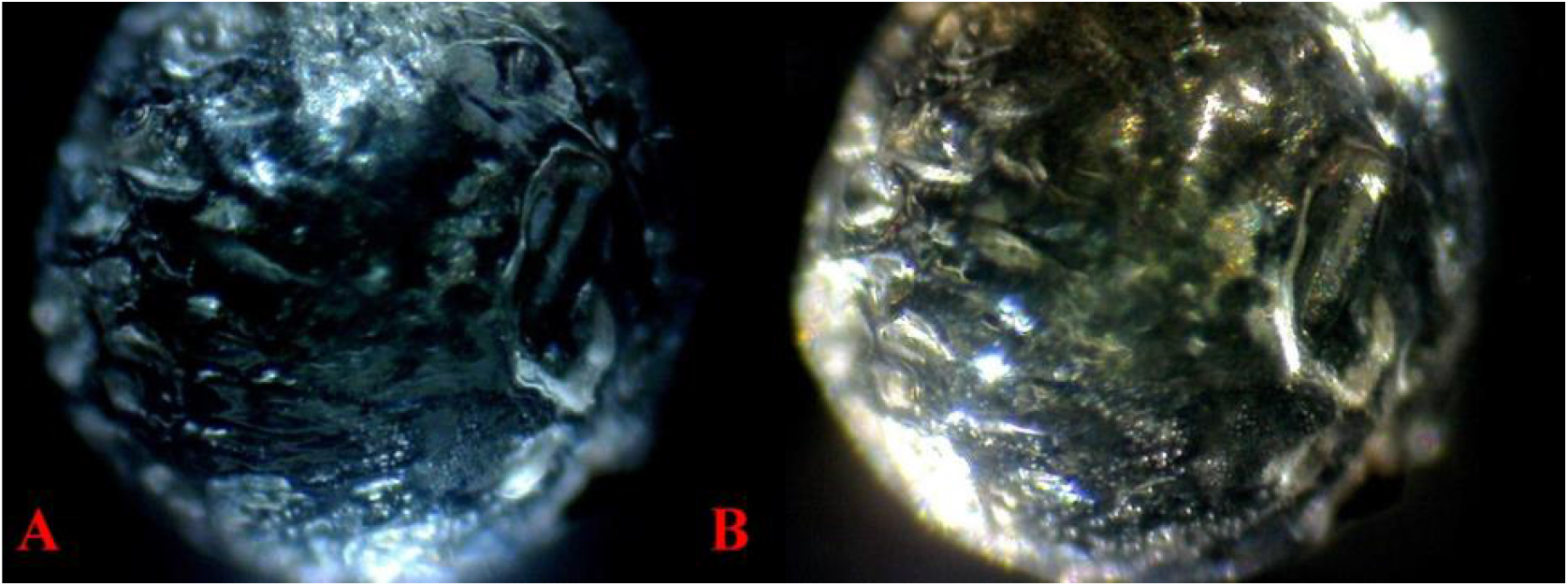
**(A)** 30 40× 14 **Pb/Pb* (full dissection and extraction*) reflected light. **(B)** The same field, reflected light with white balance adjusted.

Several lenses were intentionally crushed with compression between the glass slide and the cover slip. A clear distinction was revealed between ruptured tissue fragments (with chromatophore populations) and the fractured rigid crystalline epithelial cells (Fig 30A-B). Further distinction was visible between fractured epithelial cells forced into underlying reflective crystals within the cortex and the lens itself. Reflective qualities, both yellow and blue, were detected in epithelial cells from the germinative zone of the lens (Fig 31).

**Fig 31.**
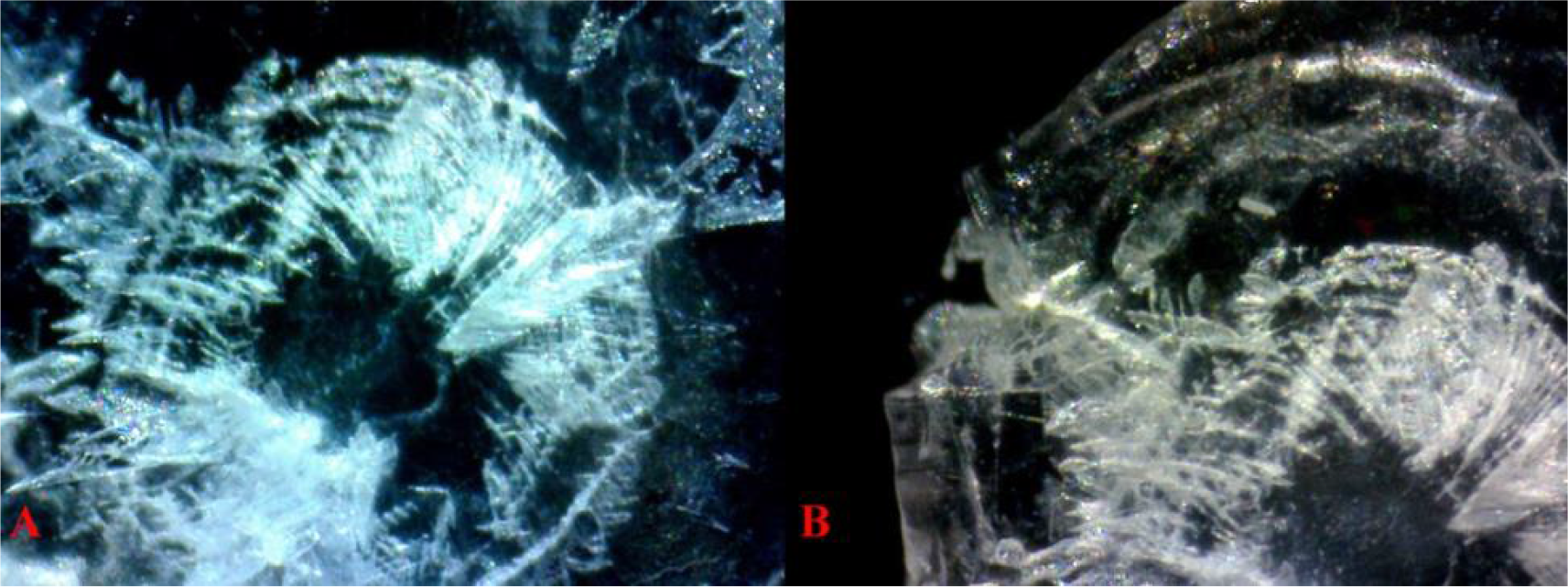
Compressed Iris. **(A)** 30 Pb 40× 24 *Pb/- (full dissection and extraction*) reflected light. **(B)** 30 Pb 40× 27 *Pb/- (full dissection and extraction*) reflected light. With white balance adjusted. In both images (top right) the ruptured surrounding membrane of the lens showing high levels violet-blue iridophores and xantho-erythrophores is present. Also, a section of membrane is visible behind the center nucleus. Scattered collections also appear in fragments of the menbrane. Fractured crystalline epithelial cells appearing as long-linear structures appear devoid of any chromatophore populations.

**Fig 32.**
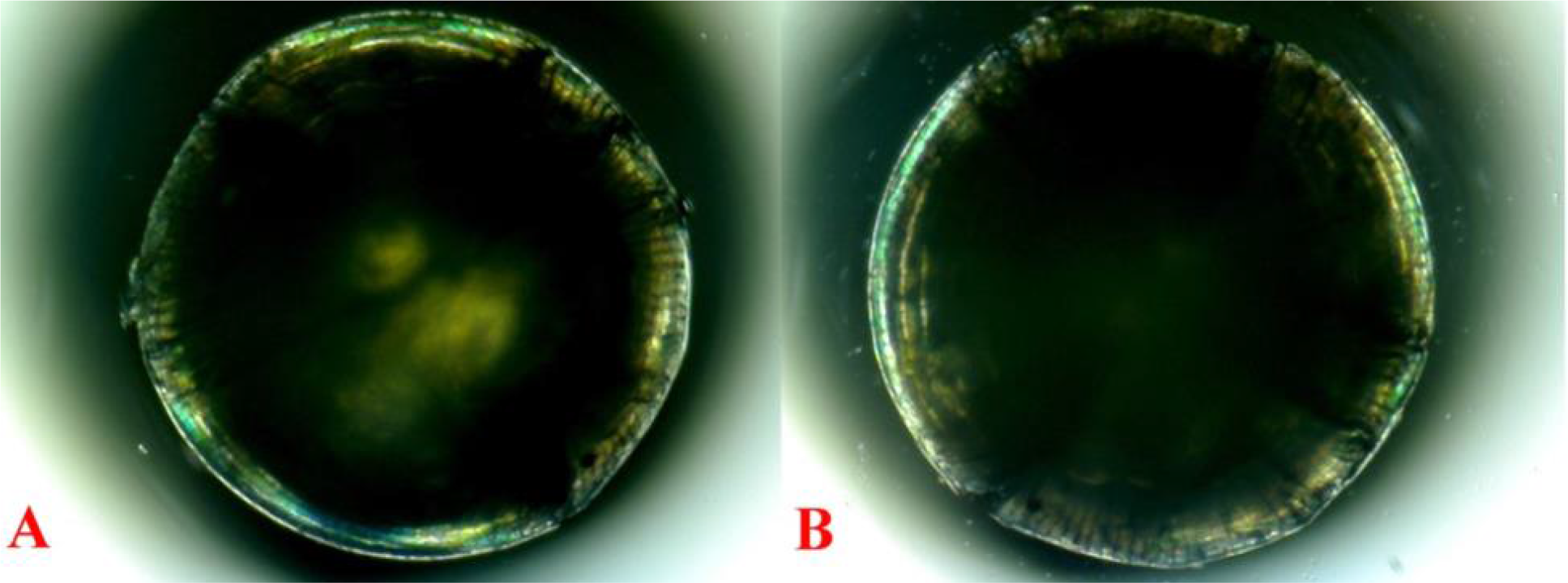
**(A)** 31 40× 27 **Pb/Pb* (full dissection and extraction*) reflected and transmitted light. **(B)** 31 40× 29 **Pb/Pb* (full dissection and extraction*) reflected and transmitted light. Reflective xanthophores and iridophores present in epithelial cells along the outer germinative zone of the lens. They are in turn reflected by the central nucleus of the lens.

**Fig 33.**
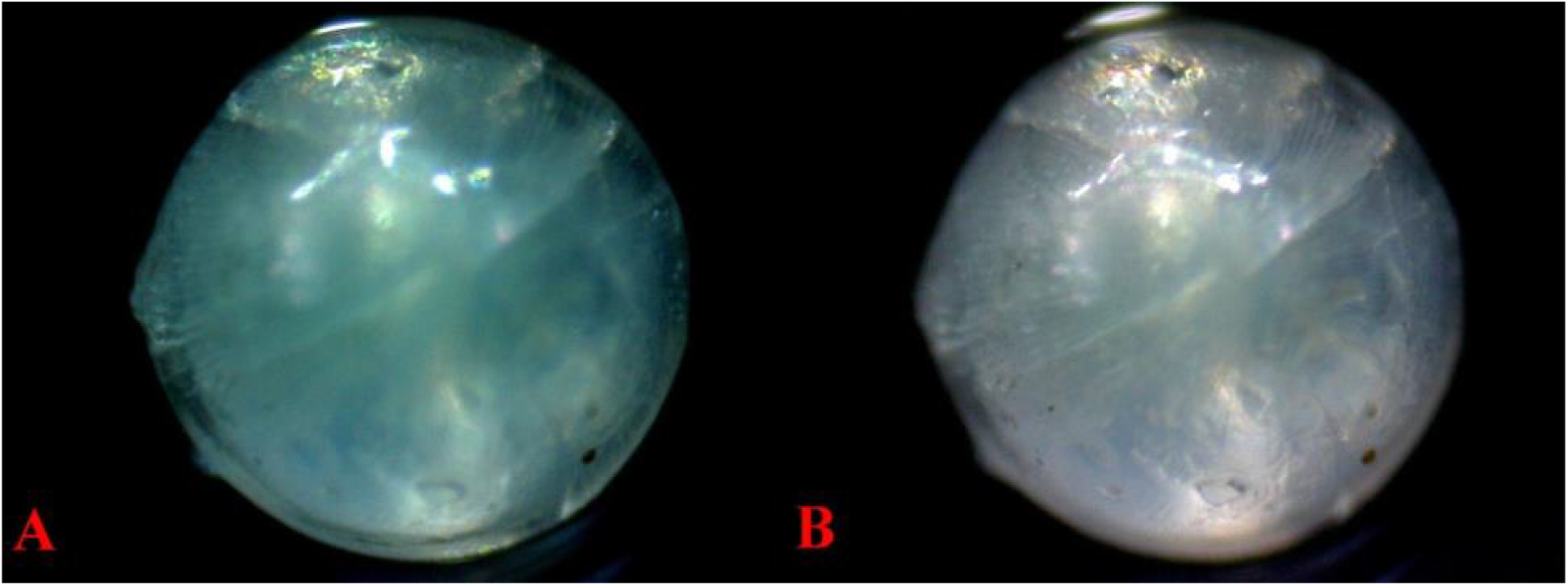
**(A)** 31 40× 27 **Pb/Pb* (full dissection and extraction*) reflected light. **(B)** The same field reflected light and white balance adjusted. Reflective pigments from the germinative zone are visible in the upper center. This lens was extracted from an aged female. Large opaque cortical cataracts “spokes” are visible in the central cortex of the lens. An apparent opaque nuclear cataract “cloudiness” appears in the central nucleus of the lens and anterior membranous inclusion is visible in the lower right. These pathological changes are consistent with UVB damage.

**Fig 34.**
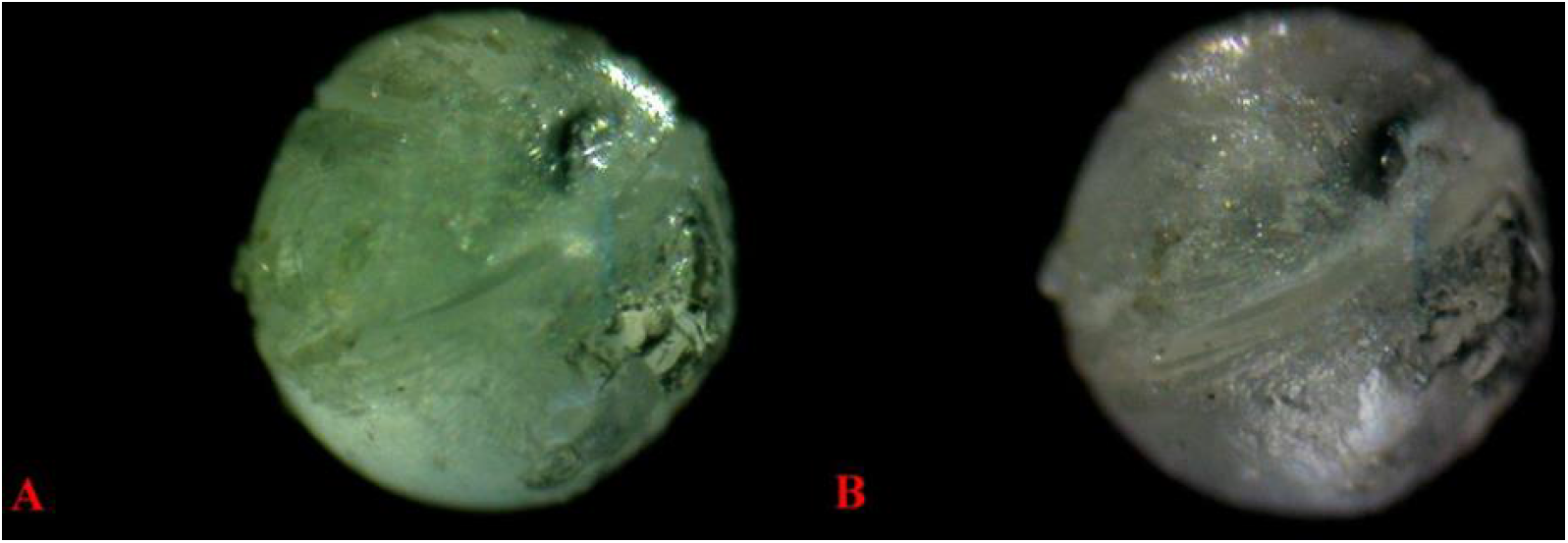
**(A)** 31 40× 27 **Pb/Pb* (full dissection and extraction*) reflected light. **(B)** The same field reflected light and white balance adjusted. Chromatophores are detected within the surrounding membrane “capsule” of the lens. The deep circumventing indentation around the lens is likely the result of underlying cortical cataracts noted previously. Large areas of melanophore and violet-blue iridophore on the lens anterior capsule (lower right) appear to result from the iris adhering to the lens (synechia).

**Fig 35.**
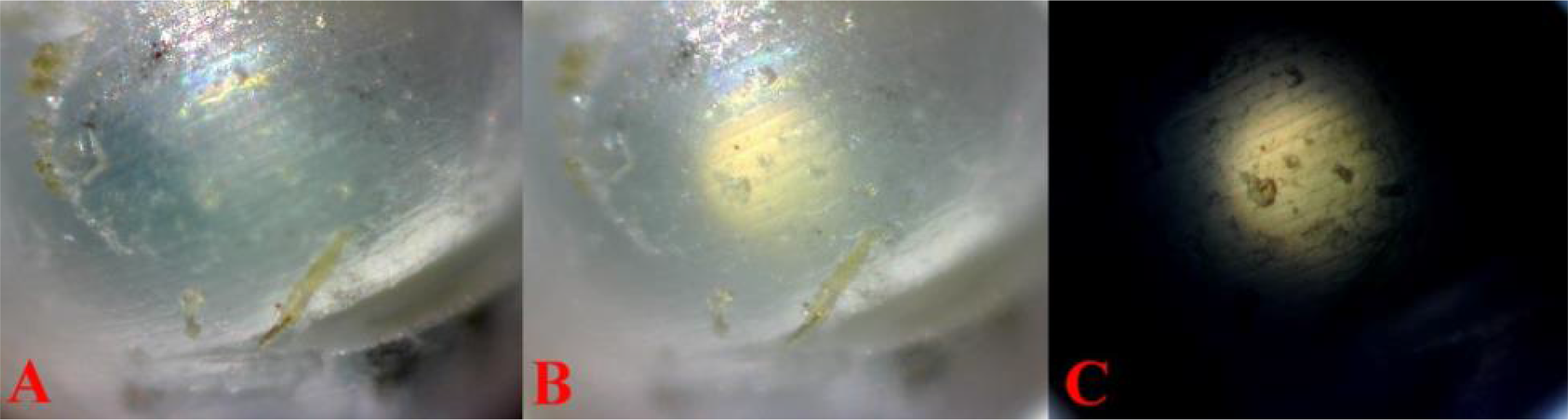
**(A)** 31 100× 2 **Pb/Pb* (full dissection and extraction*) reflected light. Isolated reflective cells detected in the surrounding membrane. Defective anterior membrane inclusions (xanthophores and melanophores were described previously). Posterior sub-capsular cataracts “granular deposits” appearing milky white. **(B)** The same field reflected and transmitted light. Defective anterior membrane inclusions (xanthophores and melanophores described previously). Posterior sub-capsular cataracts “granular deposits” appearing milky white. **(C)** The same field transmitted light. Posterior sub-capsular cataracts “granular deposits” appearing dark colored. These pathological changes are consistent with UVB damage.

**Fig 36.**
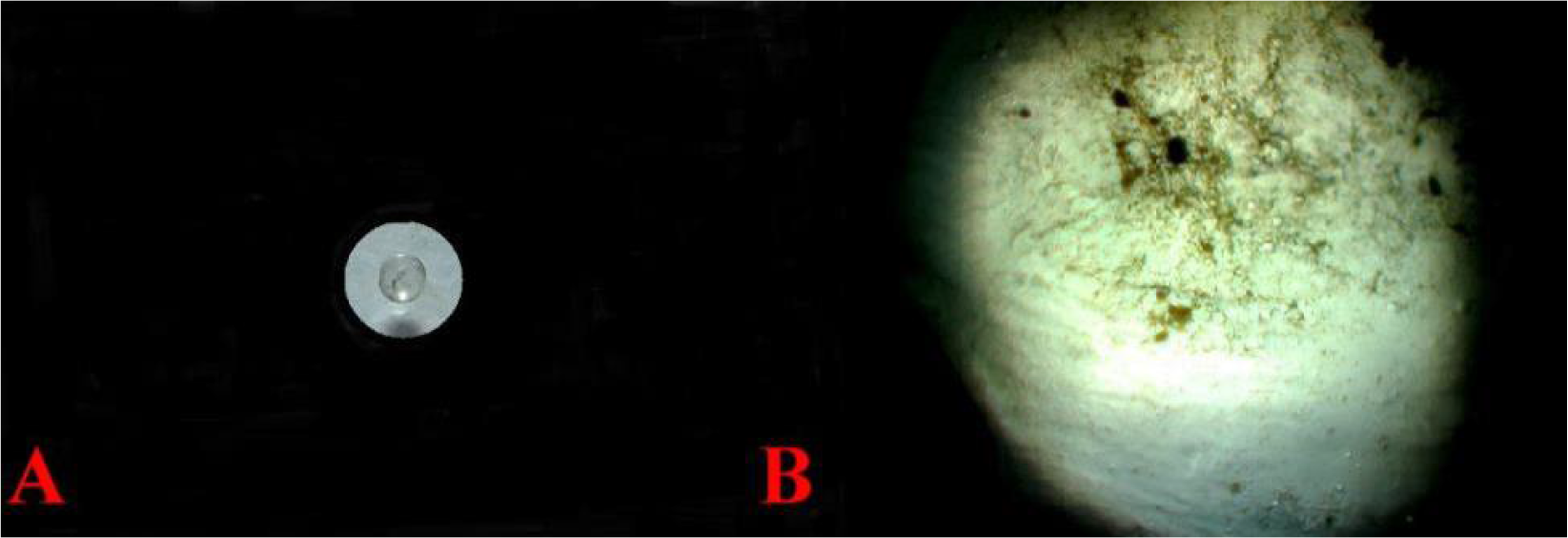
Macroscopic view of an extracted lens after multiple saline and alcohol rinses with prolonged soaking time. **(A)** 35mm, Focal length 30.3mm, F stop: F/5, Exposure time: 1/240 second, Metering mode: spot, Exposure compensation: -2step. *Pb/- (dissected*) Slave flash. Large anterior capsule inclusion is visible in the center of the lens. **(B)** 30 100× 4 *Pb/- (dissected*) transmitted light white balance adjusted. Large areas of melanophores on the lens anterior capsule (lower right) appear to result from the iris adhering to the lens (synechia).

## Discussion and Conclusions

Pb has been identified as the first polymorphic autosomal gene to be described as existent in high frequencies in wild, feral and Domestic Guppy populations. It is capable of pleiotropic effects on all existing color and pattern elements at multiple loci. It should therefore be considered a strong candidate for further studies involving “the relationships between spectral and ultrastructure characteristics” in orange ornamentation, and extending to color and/or pattern as a whole as suggested by Kottler (2014). A mechanism was identified by which Pb is capable of balancing overall color and pattern polymorphisms, in turn providing fitness through heterozygosity in diverse complex habitats (Bias and Squire 2017a, 2017b). We hope that Purple will be mapped to its linkage group.

All major classes of chromatophores (melanophores, xanthophores, erythrophores, violet-blue iridophores) and crystalline platelets in the iris and ocular media (cornea, aqueous humor, vitreous humor, outer lens membrane), and possibly the lens itself of *Poecilia reticulat*a affect the transmission of light into the eye and along the visual path to the retina. The relative proportions of these chromatophores depend upon the genotype. Thus in the case of the Metal Gold (*Mg*) mutant shown in **Fig. 28**, there is a relative increase in the number of xanthophores in the aqueous and vitreous humors. An albino mutant (not used in this study, but shown and discussed in Bias and Squire (2017c) would lack all melanophores in the ocular filter system.

Although 10 opsins were found in guppies by Laver (2011), only four opsins were expressed at high frequencies (% relative abundance): A180 ∼20% in adults, RH-2 >40% in females and juveniles, ∼30% in males, and SWS2B >25% in adults. Their expression data led them to assign the 359/389 nm to UVA SWS1, the 408nm peak to violet SWS2B, the 465nm peak (Middle Wave Length, or blue) to RH2-2, and the cones with 533nm, 543nm and 573nm (Long Wave Length or green to yellow) peaks to A180 and others at low frequencies. These are peak values, and each would produce a curve extending in both directions. They found the S180 gene produces Long Wave Length 560nm cones that would detect yellow, and the S180r gene (functioning at lower levels) produces 572nm cones, which would also respond to yellow. They appear to be the cones responding to orange and red colors. The other opsins that are present at low levels are also functionally important, especially with regard to this study. This was also reported in the Cumaná guppy by Endler (2001), and Watson (2010). Watson quotes Archer (1990) as saying that the absorption spectra for the cones of the Trinidadian guppy are somewhat different, with a UV peak at 389 instead of 359.

Grether (2008) showed the absorbance spectra for guppy carotenoids from eggs and skin. The curves were slightly different, but both peaked at around 440nm with a second lower peak at 475nm dropping sharply before 500nm. The curve continued into the lower range with a low at around 355nm and then rising again to 300nm where the graph stopped. Grether (2001) showed both the absorbance spectra for carotenoids and drosopterin extracted from orange spots of male guppies in their Fig. 3. They also showed the simulated reflectance spectra for different ratios of carotenoid and drosopterin ratios. In their Fig. 4 they showed (in addition to other values) the mean orange-spot reflectance spectra for guppies in the field. This reflectance begins at a value of “0.20” and starts dropping at around 570nm and by around 510nm has reached a low value of around “0.06”. The reflectance value of drosopterin starts dropping as its absorbance value increases. The absorbance value of guppy carotenoid starts increasing at lower wavelengths.

Actual reflectance spots showed the highest values for violet spots (Kemp 2009) in male guppies from the transplanted Trinidadian colony in an upper stream of the El Cedro River where the main predator is a Pike Cichlid, *Crenicichla alta*. “Reflectance spectra captured from the four most common colour pattern elements of El Cedro fish (orange spots, iridescent blue spots, blue/violet iridescence and iridescent green/blue) are represented… All colour elements are characterized by a strong reflectance peak…in the region of 370–390 nm, thus indicating a strong UV component. In perceptual terms, the largest and most consistent difference between the populations is that all viewers would perceive brighter iridescent blue/violet markings in introduction site fish…” Therefore these male guppies were utilizing UV light in the near visible end of the UV spectrum.

According to Weadick (2012), this pike cichlid (known both as *C. alta* and as *C. frenata*) has apparent absence of an SWS1 opsin and some synthesis of SWS2a and SWS2b UV sensitive opsins. These later two opsins may be present in low amounts and the authors state that “this fish may be relatively insensitive to UV light and unable to discriminate hues in the lower part of the visual spectrum. If this is the case, guppies could potentially use UV light as a private communication channel as they do possess an UV-sensitive cone”. They refer to Archer (1990). They also point out that there is some variation in the visual capability of Pike Cichlids from different sites, and it is quite possible that other populations may have completely lost the ability to see in the UV spectra, as had been earlier stated by Endler (1991).

To summarize our comments so far then: guppies produce opsins that are sensitive to UV down to at least 250nm and probably somewhat lower. This range includes the reflectance peaks in the region of 370-390nm. As would be expected, guppies are able to detect the UV reflectance of their neighbors, as well as colors in the visible spectrum. An obvious question then would be “Why are the eye capsule, cornea, aqueous humor, vitreous humor, outer lens membrane pigmented? What are the selective benefits provided by this pigmentation?” Ultra violet light includes the range of 100 - 400nm. It is destructive; its absorption causes cell damage and cell death.

Melanin is known to absorb UV light and thus prevent it from harming cells. Human dark skin color and tanning are examples of this protective function in humans (McGraw 2005). Further, a number of reports indicate that both the cornea and the lens act as UV filters in many if not most species of vertebrates (Nelson 2001). It is generally held that when UV filtration occurs the cornea is the first UV filter, and the lens is the second. Thorpe (1993) found that the guppy cornea (of an unidentified population) transmits 50% of incident UV at 315nm. This indicates that the guppy lens is not a major filter of UVA wavelengths (320-400nm). But it may be a significant blocker of UVB rays (280-320nm).

Judging from Fig. 3 of Grether (2001) the absorbance values of drosopterin become significant by 525nm and increases to a peak around 480nm and gradually diminishing. Likewise the absorbance values for guppy carotenoids seem to become significant around 490nm and extend to below 400nm (the limit of their figure). They point out that absorbance values for intact cells will vary from those of the extracts used in their study. Carotenoids are antioxidents (Svensso 2011). Thus melanin, drosopterin and guppy carotenoids absorb UV light and can provide protection against its damage.

The presence of melanophores (melanin), xanthophores (carotenoids), xantho-erythophores and iridophores (guanine absorbs UV light) all provide protection from UV-induced damage to the structures over which they are located. These cells are present in a very thin layer over the cornea and the lens, and thus would allow the passage of UV and other wavelengths through them with minimal absorption. This would cause accumulated damage over time and would produce the cataracts and milky deposits seen in the lens from an old female in **Fig. 32 **and** 34.** The thick layer of melanophores and the dense layer of violet-blue iridophores in the iris provide protection against UV-induced damage.

What is the function of the violet-blue iridophores and xantho-erythrophores found in the aqueous and vitreous humors? These humors are part of the pathway of light that passes through the cornea, the aqueous humor, the lens, the vitreous humor, and then stimulates receptor cells in the retina itself. The aqueous humor is interposed between the cornea and the lens and the vitreous humor between the lens and the retina. As such they may act as visual filters, perhaps by absorbing additional UV light, and they attenuate the degree of exposure of the retinal cells themselves. A number of UV-absorbing compounds are known (proteins, amino acids and derivatives thereof, ascorbic and uric acid) from cornea, aqueous humor, and lens of vertebrate eyes (Ringvold 2003), but we are unaware of any reported ocular filter systems using intact pigment cells rather than dissolved molecules. (We did not examine dissolved molecules from the eyes in this study.) These cells have a greater absorptive ability for UV in the wavelengths below those used by the guppy opsins. Melanins, carotenoids and pteridines are also antioxidants and would tend to remove free radicals before they harm important cells and structures.

The medical role of pterins (pteridines) as antioxidants in immunology is so important that pterin levels are used as a clinical biomarker of immune performance (McGraw 2005). UV light has more energy per photon than any other wavelength of light that reaches the earth’s surface. These highly energetic photons damage many kinds of biological molecules, of which DNA and proteins are the most obvious. They also cause chemical reactions that produce “reactive oxygen species” (ROS). These chemically reactive molecules then cause additional damage. Dissolved organic carbon (DOC) reduced UV penetration and thus is a protective factor in the aquatic environment. Fish in shallow water are more at risk of UV damage. This damage may affect fish eggs and young fish as well as adults. (Zagarese, 2001; Gouveia 2015). When studying Atlantic cod eggs, Kouwenberg (1999), found no evidence of detrimental effect of UVA radiation (320-400nm). However they found considerable mortality from UVB (280-320nm). Douglas (1999) states that “no ocular structure will transmit significant amounts of radiation below about 310nm due to absorption by its nucleic acids and various structural protein components, such as aromatic amino acids.” Of course this absorption damages the molecules and structures involved.

This does not infer that UVA is not potentially harmful to guppies! All UV radiation is potentially harmful. In this regard, the melanophores, xanthophores and iridophores reported in the membranes surrounding the spinal cord of guppies (Bias and Squire, 2017b) may also provide protection against UV damage (Gibson 2009). Thus the potentially less harmful UVA is being used in the guppy UV ornament-UV opsin system. But the more harmful UVB needs to be filtered out more thoroughly. It would seem that the tendency of young guppies to hide in dense plant growth may shield them from UV damage as well as provide a hiding place from predators. It is conceivable that the violet-blue iridophores (absorbance of UVB and potential scattering of UVA) and xantho-erythrophores (absorbance) have a strong filtering capacity in the UVB range.

Douglas (1989) comments that since fish with short-wave absorbing filters deprive themselves of the potential benefits of UV sensing they must gain a sufficient adaptive advantage to make up for it. But we would suggest that these do not need to be mutually exclusive alternatives. They refer to Walls’ (1933) proposal that “short-wave absorbing filters may lead to increased visual resolution, contrast and visual range, by decreasing (i) the degree of chromatic aberration, (ii) the amount of scattered light reaching the retina, and (iii) the glare from the bright down welling illumination, all of which are highest at short wavelengths.” We propose that the guppy has achieved a balanced system in which a reduced amount of UVA reaches the retinal UV receptors, while at the same time excluding much of the damaging UVB radiation.

The presence of large numbers of iridophores in this system is noteworthy. Violet-blue iridophores were present in the cornea, the outer lens membrane was saturated with violet-blue iridophores, and they were present in high numbers in both the aqueous and vitreous humors. Tovée (1995) suggests a different possible aspect to UV sensitive cones in animals like the guppy. He suggests that the UV-sensitive cone density is not sufficient to provide a high-resolution system on its own. The responses of UV cone receptors may have to be pooled with the results of other types of cones in order to provide a higher order integrated sensory “image” at the level of the brain. While he then argues against his own suggestion, it may be worth reconsidering.

A final point should be re-emphasized. Although there are numerous pigmented cells in the direct light pathway from the cornea to the retina, and these cells are expected to filter out considerable amounts of UV light, especially UVB, these cells do not prevent sufficient UVA from reaching the retina and stimulating the UV cones therein.

## Photo Imaging

Photos by the senior author were taken with a Fujifilm FinePix HS25EXR; settings Macro, AF: center, Auto Focus: continuous, varying Exposure Compensation, Image Size 16:9, Image Quality: Fine, ISO: 200, Film Simulation: Astia/Soft, White Balance: 0, Tone: STD, Dynamic Range: 200, Sharpness: STD, Noise Reduction: High, Intelligent Sharpness: On. Lens: Fujinon 30x Optical Zoom. Flash: External mounted EF-42 Slave Flash; settings at EV: 0.0, 35mm, PR1/1, Flash: −2/3. Photos cropped or brightness adjusted when needed with Microsoft Office 2010 Picture Manager and Adobe Photoshop CS5. All photos by the senior author.

## Microscopy

All Digital Image processing by conventional bright and dark field equipment. AmScope M158C. Camera(s): 1. MD35, Resolution: 0.3MP 2. MD200, Resolution: 2MP USB Digital, Sensor: Aptina (Color), Sensor Type: CMOS. Software: AmScope for Windows. An attempt was made to restrict ambient light during both daytime and nighttime imaging of specimens. Imaging was performed with reflected or transmitted practical light sources as indicated. Where delineation in results warranted, a series of three photos from each location were taken and presented in the results; reflected (top light only), transmitted (bottom light only), combined reflected + transmitted (top and bottom light).

For purposes of this study low resolution photos were often preferred over higher resolution for clarity at settings of 40×, 100X or 400X. No images were stained. As identified, individual images are full body (non-dissected), or manually de-fleshed (dissected) skin samples. Samples were air dried for minimal time periods of less than one hour for aid in dissection. All samples and images from right side of body, unless otherwise noted. No cover glass was utilized, to reduce damage to chromatophore shape, structure and positioning. No preservatives were used during imaging, though rehydration was done as needed for clarity. All photos were by the senior author.

## Ethics Statement

This study adhered to established ethical practices under AVMA Guidelines for the Euthanasia of Animals: 2013 Edition, S6.2.2 Physical Methods (6).

All euthanized specimens were photographed immediately, or as soon as possible, after temperature reduction (rapid chilling) in water (H20) at temperatures just above freezing (0°C) to avoid potential damage to tissue and chromatophores, while preserving maximum expression of motile xantho-erythrophores in Pb and non-Pb specimens. All anesthetized specimens were photographed immediately after short-term immersion in a mixture of 50% aged tank water (H_2_0) and 50% carbonated water (H_2_CO_3_).

All dried specimens photographed immediately after rehydration in cold water (H20). Prior euthanasia was by cold water (H20) immersion at temperatures just above freezing (0 °C). MS-222 (Tricaine methanesulfonate) was not used to avoid the potential for reported damage and/or alterations to chromatophores, in particular melanophores, prior to slide preparation.

## Competing Interests and Funding

The authors declare that they have no competing interests. Senior author is a member of the Editorial Board for Poeciliid Research; International Journal of the Bioflux Society, and requested non-affiliated independent peer review volunteers.

The authors received no funding for this work.

## Acknowledgements

To my best friend and wife Deana Bias, for her support and persistence over the last several years in this four part study… To my co-author and dear friend Rick Squire for his patience as a mentor… To those Domestic Breeders who willingly provided additionally needed pedigree strains and study populations for completion of this paper…

## Supporting Information

S1 Materials; Slide Specimen Photos

## Notes

This publication is number three (3) of four (4) by Bias and Squire in the study of Purple Body (*Pb*) in *Poecilia reticulata*:

1. The Cellular Expression and Genetics of an Established Polymorphism in *Poecilia reticulata;* “Purple Body, (*Pb)”* is an Autosomal Dominant Gene,

2. The Cellular Expression and Genetics of Purple Body (*Pb*) in *Poecilia reticulata*, and its Interactions with Asian Blau (*Ab*) and Blond (*bb*) under Reflected and Transmitted Light,

3. The Cellular Expression and Genetics of Purple Body (*Pb*) in the Ocular Media of the Guppy *Poecilia reticulata*,

4. The Phenotypic Expression of Purple Body (*Pb*) in Domestic Guppy Strains of *Poecilia reticulata*.

